# Speed-dependent place- and time-field shifts do not require explicit temporal coding

**DOI:** 10.64898/2026.07.17.739217

**Authors:** Federico Szmidt, Camilo J. Mininni

## Abstract

Place and time cells are widely thought to provide complementary representations of spatial location and elapsed time in the hippocampus. Recent experiments reported CA1 neurons whose place and time fields shift systematically with running speed, suggesting that representations of space and time are integrated and compete within a common neural code. Here we show that speed-dependent place- and time-field shifts can arise without explicit temporal coding. Using a hierarchy of computational models, we demonstrate that these phenomena emerge when the velocity of the internal estimate of position becomes progressively less sensitive to increases in the animal’s speed. Recurrent neural networks trained exclusively for path integration spontaneously developed this behavior. Their analysis revealed a circuit mechanism in which weakly direction-selective neurons stabilize the activity bump at low speeds and progressively release this brake as speed increases. Finally, recurrent networks trained to jointly encode position and elapsed time distinguished genuine spatiotemporal representations from apparent temporal tuning generated by the task-induced correlation between space and time. Together, our results provide an alternative interpretation of speed-dependent place- and time-field shifts, identify a computational mechanism that extends continuous bump attractor models, and generate experimentally testable predictions.

## 1 Introduction

The ability to represent space and time is fundamental to animal behavior. Accurate estimates of position and elapsed time are required for navigation, the organization of actions, and the formation of episodic memories [1, 2]. A large body of work has implicated the hippocampal-entorhinal system in these computations [2, 3]. Place cells [4], grid cells [5], head-direction cells [6] and other spatially tuned neurons [7, 8] provide complementary representations of location and orientation, while ramping activity [9] and time cells [10–13] have been proposed to encode the temporal structure of behavioral episodes.

Particular attention has been devoted to hippocampal time cells, neurons that fire selectively at specific moments following a salient event or cue [10, 11]; [12]. Time-cell activity has been interpreted as a neural substrate for the temporal organization of memory, complementing the spatial representation provided by place cells [2, 14]. However, the interpretation of time-selective firing remains challenging because elapsed time is often strongly correlated with other behavioral variables, including traveled distance, position, speed, sensory input, motor state and task phase [15–17]. Consequently, apparent temporal coding can arise from mechanisms that do not explicitly represent time [15–17]. Previous studies have shown, for example, that path-integrated distance or traveled distance can generate neural responses that resemble temporal tuning when behavioral variables are constrained [15, 17]. An additional complication is that neurons frequently exhibit mixed selectivity. Many hippocampal neurons are tuned not only to elapsed time but also to spatial location [15, 18], and mixed representations of space and time have been reported across multiple brain regions and behavioral paradigms [19–21]. Such mixed tuning is often interpreted as evidence for a joint spatiotemporal code [2, 18, 21]. Yet the mechanisms that generate these responses remain poorly understood. In particular, it remains unclear to what extent mixed selectivity reflects an explicit representation of time, as opposed to the interaction between spatial coding and behavioral variables that co-vary with time [15, 16, 18].

Recent work has brought this question into sharper focus. Chen et al. [18] recorded CA1 neurons while mice navigated one-dimensional virtual and physical tracks and reported a substantial population of neurons with mixed selectivity for space and time. Remarkably, the spatial and temporal firing fields of these neurons changed systematically with running speed. As speed increased, spatial fields shifted towards more distant locations along the track, whereas temporal fields shifted towards shorter elapsed times from trial onset. At the population level, neurons also exhibited an apparent trade-off between spatial and temporal coding, such that stronger encoding of one variable was associated with weaker encoding of the other. These findings were interpreted as evidence that spatial and temporal representations are integrated and compete within a common neural code. Whether such effects necessarily require an explicit space-time representation, however, remains unresolved. In the tasks used by Chen et al., animals moved along one-dimensional trajectories with a fixed direction of travel, creating strong correlations between position, elapsed time and speed. Under such conditions, changes in spatial coding across behavioral states may generate apparent temporal effects, even in the absence of an independent temporal code [15, 17, 22, 23]. This raises a fundamental question: do the reported field shifts and apparent space-time competition require a genuine temporal representation, or can they emerge from a neural system that encodes space alone?

Here we address this question using a hierarchy of computational models. We first show that the behavioral correlation between position and elapsed time causes pure place and time cells to exhibit apparent field shifts. We then analyze a continuous line-attractor model that, by construction, encodes position but not elapsed time. We show that speed-dependent field shifts and apparent space-time competition emerge naturally when attractor speed becomes progressively less sensitive to animal speed, corresponding to a concave speed-response function. We next show that the same phenomena emerge in recurrent neural networks trained exclusively for spatial encoding. Analysis of these networks revealed a simple brake mechanism, in which weakly direction-selective neurons progressively lose influence as speed increases, producing the concave speed response. Finally, we trained recurrent networks to jointly encode space and time and used them to distinguish genuine from apparent spatiotemporal representations. Together, our results suggest that the key observations reported by Chen et al. may reflect a previously unrecognized property of hippocampal spatial representations: a concave relationship between the speed of the internal position estimate and the animal’s speed.

## 2 Results

### 2.1 Pure place or time cells cannot explain simultaneous spatial and temporal field shifts

We began by asking whether the key observations reported by Chen et al. necessarily require neurons that jointly encode position and elapsed time, or whether they could instead arise from simpler mechanisms. As a first step, we analyzed the behavior expected from idealized pure place cells and pure time cells in the same behavioral paradigm. Although these cells represent only one variable, the structure of the task itself correlates position and elapsed time, raising the possibility that apparent place or time fields could emerge in the complementary reference frame. To that end, we consider an animal running on a linear track (schematized in Fig. 1a). The leftmost position is taken as the spatial origin, time is measured from the beginning of an episode and the animal runs from left to right. In this setting, a pure place cell fires at a specific location on the track, regardless of the animal’s speed (Fig. 1b). When its firing field is represented in the temporal reference frame, however, its apparent time field shifts toward shorter elapsed times as speed increases (Fig. 1c). Conversely, when the firing field of a pure time cell is represented in the spatial reference frame, its apparent place field shifts forward along the track (Fig. 1d), while its time field remains centered at a fixed elapsed time regardless of speed (Fig. 1e).

**Figure 1:**
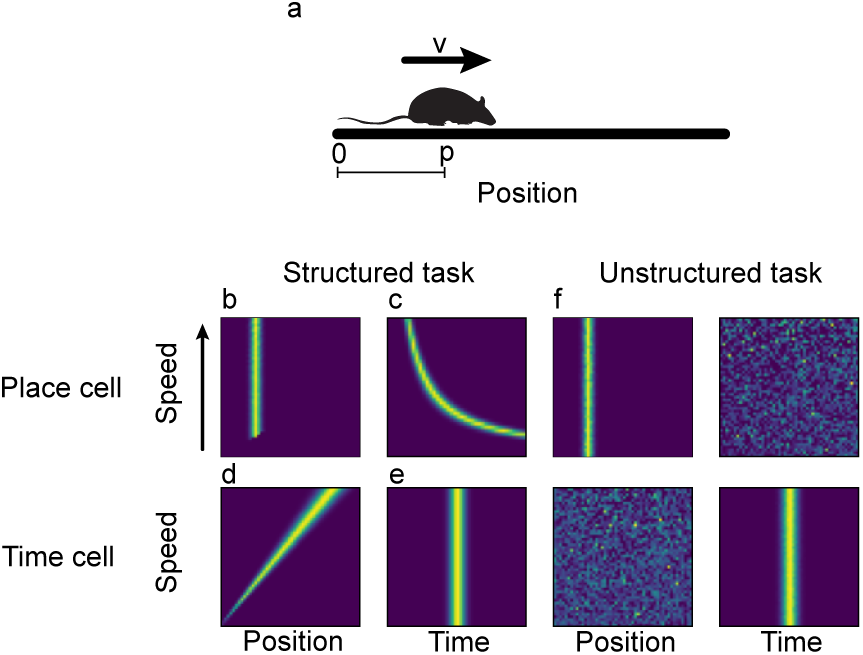
Pure place cells or time cells cannot explain simultaneous place and time field shifts. **a** A simulated animal walks along a linear track with constant speed v, and occupies position p at time t. **b** Speed-ordered spatial rate map for a simulated pure-place cell shows no shift of the place field. **c** Speed-ordered time rate map for a simulated pure place cell shows a backward time field shift. **d** Speed-ordered spatial rate-map for a simulated pure time cell shows the place field shifting forward. **e** Speed-ordered time rate-map plot of a pure time cell shows no shift of the time field. **f** In the unstructured task each direction is chosen uniformly at random, thereby decorrelating space and time; place cells show no encoding of time, and time cells show no encoding of space.

These shifts arise solely from the behavioral correlation between position and elapsed time imposed by the task. For an animal moving at constant speed *v* = *p/t* (*p* being the animal’s position) then a pure place cell will have an apparent time field at time *p_n_/v*, where *p_n_* is the animal’s position when the neuron activates. This time field recedes as speed increases. In contrast, a pure time cell will fire at position *t_n_v*, where *t_n_* is the time at which the time cell activates, and the apparent place field will move forward as the speed increases. When the animal instead moves from uniformly sampled initial positions and movement directions, the correlation between position and elapsed time is removed and these apparent shifts disappear (Fig. 1f). Thus, even pure place cells and pure time cells can exhibit apparent field shifts and space-time trade-off in a 1D track when movement in only one direction is allowed (a *structured task*), while the shifts disappear when the space-time correlation induced by the task is removed (an *unstructured task*). Nevertheless, this simple model cannot explain the observations of Chen et al., in which individual neurons simultaneously exhibit both place and time fields that shift with running speed.

### 2.2 Field shifts and space-time trade-off emerge in a continuous line attractor with concave speed response

To further explain the potential space-time coding interaction and its alternative explanations, we implemented a simple continuous line attractor, designed to only integrate the animal’s speed and therefore lacking genuine time-encoding neurons. The model is characterized by a single state variable *n*(*t*) and its derivative with respect to time *dn*(*t*)*/dt* (the attractor speed). We make the following assumptions: (i) each attractor position *n* corresponds to a single active neuron; (ii) the attractor speed is a function of the animal’s speed *v*(*t*); (iii) *n* is set to zero at the start of an episode; and (iv) *v*(*t*) and *dn/dt* are always positive. Consequently, the modeled animal only moves in one direction, as in the experiments of Chen et al., and the line attractor tracks its displacement. If the attractor speed scales proportionally with animal speed, that is, *dn/dt* = *v*(*t*), the attractor performs perfect path integration and each position along the track can be mapped back to an active neuron in the line attractor. In this case, neurons behave as pure place cells. The key question, however, is what happens when the attractor speed no longer scales proportionally with the animal’s speed. To address this, we assumed that the attractor speed is given by *dn/dt* = *f* (*v*), where *f* is a positive, non-decreasing, and concave function (Fig. 2a). Under this assumption, increasing the animal’s speed produces progressively smaller increments in attractor speed. As shown analytically in the Methods, this simple modification is sufficient to generate speed-dependent firing-field shifts resembling those reported by Chen et al.

**Figure 2:**
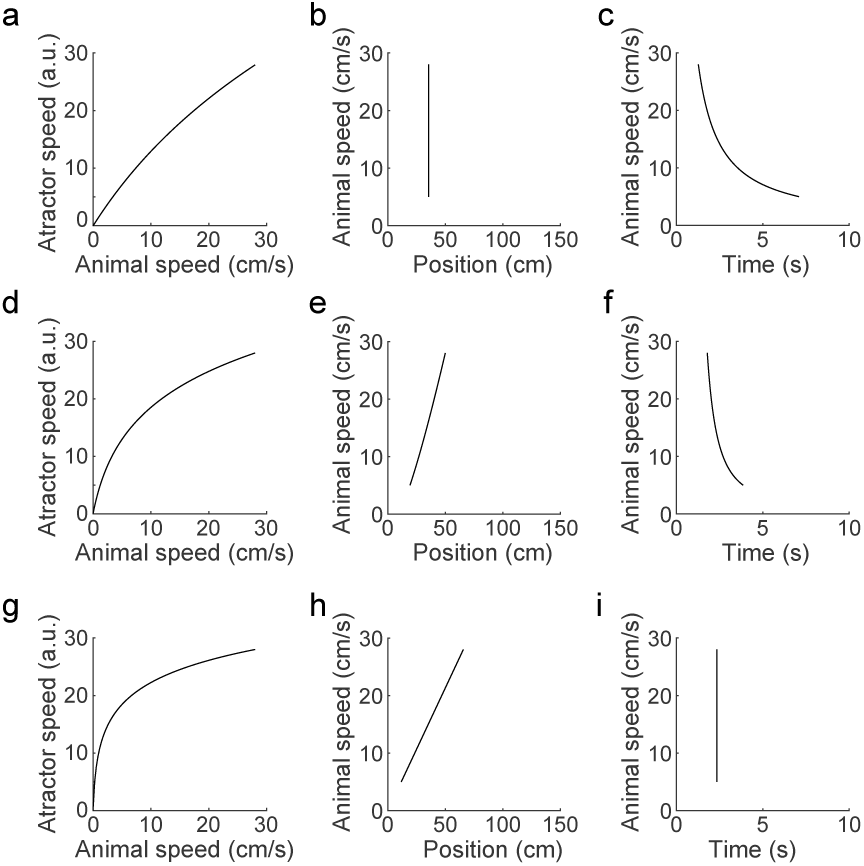
The concavity of the attractor speed-response determines field shifts. **a** Attractor speed as a function of animal speed (*v*) was defined as 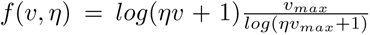, with *v_max_* = 28 cm/s. Other concave functions produce similar qualitative results. **b-c** Animal speed as a function of animal position **b**, and elapsed time since episode onset **c** for *η* = 0.05 and neuron *n* = 50. **d-f** Idem **a-c** but for *η* = 0.5, producing field shifts that closely match those shown in Figure 4 of Chen et al. **g-i** Idem **d-f** but for *η* = 5. As *η* increases, neurons tend to become more like time cells, and less like place cells. Field shifts were evaluated in the 5 - 28 cm/s velocity range, following the controlled-velocity experiments of Chen et al.

To explore the consequences of different degrees of concavity, we defined *dn/dt* = *f* (*ηv*), where the parameter *η* controls the attractor-speed response. For small values of *η*, the concavity of *f* is minimal, so the attractor behaves as a perfect integrator and neurons remain close to pure place cells (Fig. 2 a-c). Increasing *η* makes the attractor speed progressively more concave. At intermediate values, neither place nor time fields remain fixed; instead, both shift with animal speed (Fig. 2 d-f; compare with Figure 4b in Chen et al.). Finally, for large values of *η*, neurons approach pure time cells, exhibiting stable time fields and strongly shifting place fields (Fig. 2 g-i). Thus, varying a single parameter produces a continuous transition from place coding to mixed coding and finally to time coding. Moreover, the smaller the shifts become in one reference frame, the larger they become in the other, reproducing the apparent space-time trade-off reported in Figure. 6 of Chen et al.

### 2.3 Recurrent neural networks trained exclusively for spatial encoding shows shifting fields and space-time trade-offs

The above results show that a simple continuous line attractor can explain why place cells may appear to be time cells when space and time are correlated at the behavioral level. To move beyond abstract models, we trained a vanilla recurrent neural network (RNN) to solve a one-dimensional path integration task using Backpropagation Through Time [24] (Fig. 3a-b). We simulated an animal moving along a circular track and trained the network to report the sine and cosine of its angular position *θ*, as in previous studies [25]. Representing position in this way ensures that all locations on the track are encoded equivalently and avoids discontinuities at the boundaries. Importantly, the network was trained on an unstructured task, in which trajectories started from random positions, movement direction switched randomly at each time step with probability 1*/*2, and speed was independently sampled from a normal distribution. Thus, no fixed relationship between position and elapsed time was present during training. Additionally, with probability 0.1, speed was set to zero, allowing the network to learn to maintain a stable position estimate while the simulated animal remained stationary.

**Figure 3:**
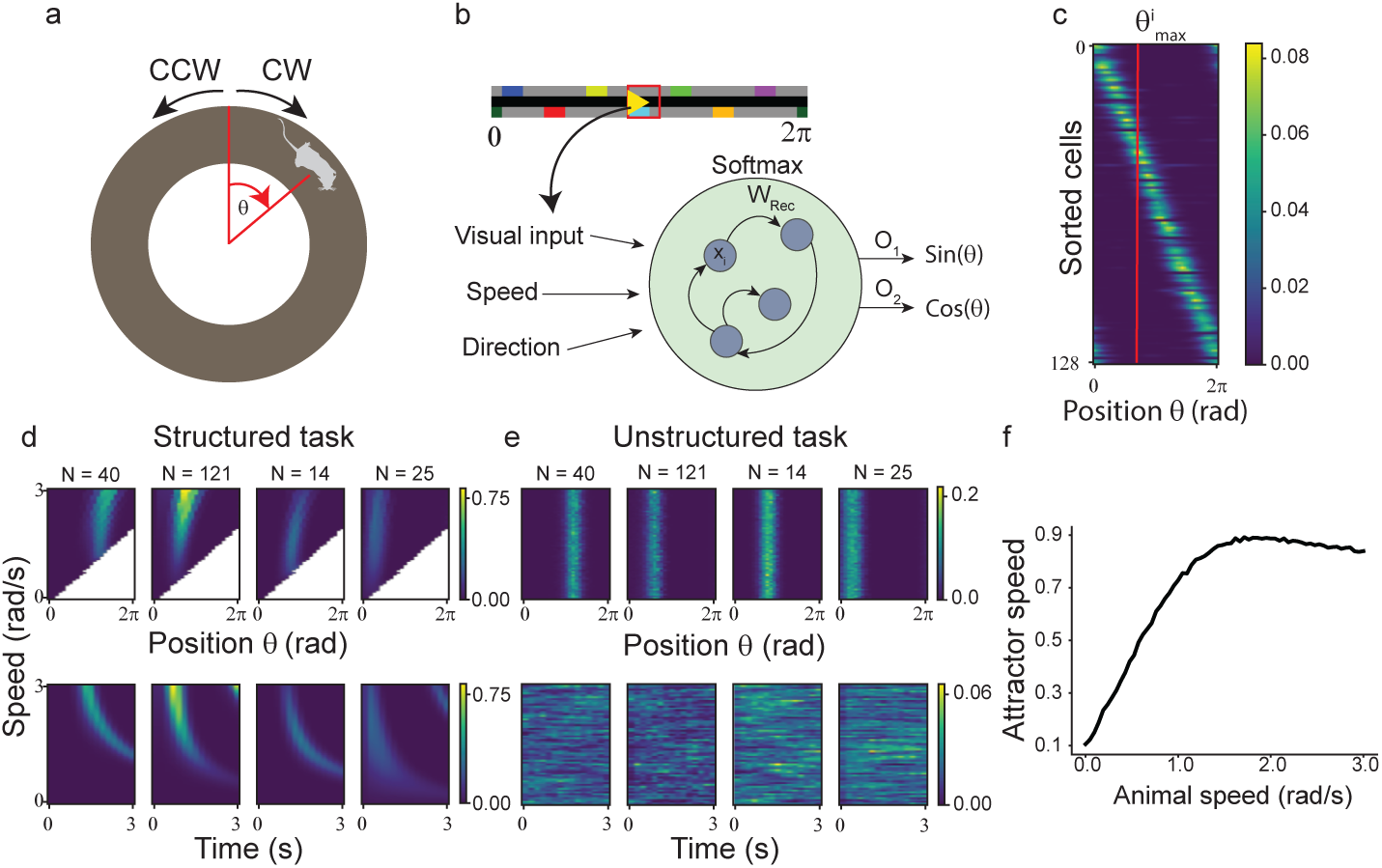
An RNN trained only for path-integration shows place- and time-field shifts. **a** Task schematic; During training, a simulated animal navigates a one-dimensional circular track with random initial positions, running speeds, and movement directions, either clockwise (CW) or counterclockwise (CCW). The angular coordinate *θ* denotes the simulated animal’s position on the track. **b** Model’s architecture (see Methods): The network receives a scalar speed input, a one-hot direction input and a visual input; recurrent weights are constrained to be positive, while the softmax activation function implements competition. *x_i_* denotes the activation of neuron *i*. **c** Rate maps of all cells on random trajectories, sorted by position of maximum firing reveal place fields that tile the entire track. 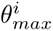 denotes the position at which cell *i* reaches its peak activation. **d** Top: Speed-ordered rate maps show place fields shifting forward in trajectories starting from the same initial position, movement direction, and fixed speed. Bottom: Speed-ordered time-averaged activity plots show a backward shift in time fields. **e** Same as (d) but for trajectories with random initial positions and directions have no place field shift, and no encoding of time. **f** Attractor speed, defined as the norm of the change in the normalized activity vector, 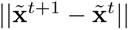 (see Methods), is a concave function of speed.

Place cells are known to update their activity using both self-motion cues and external sensory signals [26, 27]. To capture these two sources of information, the network received the animal’s speed together with a one-hot direction input, indicating whether movement was clockwise (CW) or counterclockwise (CCW), as well as a visual input presented at random times. This design ensured that the network could rely on both path integration and landmark information to estimate position. Finally, we constrained all recurrent weights to be non-negative and implemented competition among recurrent units using a Soft-max activation function. This architecture approximates a recurrent network of excitatory neurons subject to global inhibition while maintaining a sparse population code.

After training, the model successfully learned to perform path integration and approximately maintained a fixed output when the speed input was zero (Fig. S1a-c). The resulting network developed place cells that tiled the entire track (Fig. 3c) and when evaluated on trajectories with fixed speed and direction, the model exhibited a forward shift of place fields at high running speeds, resembling the results reported by Chen et al., and illustrated with four representative cells in Fig. 3d. (all cells shown in Figure S2). Likewise, aligning activity in the temporal reference frame shows time fields that shift backwards. In contrast, when we evaluated the model in trajectories with random initial positions, speeds and directions, the place fields did not shift with speed, while the time fields disappeared altogether (Fig. 3e). It is important to note that this behavior emerged in a model that was only trained to solve a spatial task, without any explicit need to encode time. These results replicate in a recurrent neural network what we previously showed in the line attractor model, and confirm that the shifts observed by Chen et al are not necessarily due to a genuine mixed encoding of space and time.

The simple line attractor model presented in the previous section suggested that firing field shifts arise from the concave relationship between animal speed and attractor speed. To determine whether the same relationship emerges in the trained recurrent network, we estimated attractor speed from the network’s population activity. Specifically, we defined the RNN attractor speed as the norm of the difference between activity vectors at two consecutive time steps, after normalizing each neuron’s activity by its maximum activation across all episodes and velocities (see Methods). Applying this measure on trajectories with fixed speed and averaging over time and random initial positions revealed that estimated attractor speed is a concave function of animal speed (Fig. 3f). This behavior is consistent with the line attractor model, and further supports the hypothesis that a concave speed response underlies the observed space and time firing-field shifts. Together, these results show that a recurrent neural network model optimized solely for spatial encoding can account for the place- and time-field shifts observed by Chen et al., and that decorrelating space and time abolishes the apparent temporal coding exhibited by the network.

### 2.4 The trained model implements a CBAN-like mechanism

So far, we have shown that the model exhibits a concave speed response capable of explaining the firing field shifts observed by Chen et al. However, the origin of this concave response within the trained network remains unclear. To uncover the underlying mechanism, we first sought to understand how the network solves the path-integration task. The emergence of place cells tiling the entire circular track suggested a resemblance to continuous bump attractor networks (CBANs), a class of models widely used to explain head-direction and place-cell systems [28–32]. Although many variants exist, one-dimensional CBANs typically comprise three distinct populations of neurons: a central ring of cells that is not direction-selective (the central cells), and two rings of clockwise (CW)-and counterclockwise (CCW)-selective cells (the directional cells). The central cells are connected through a Mexican hat connectivity profile, with excitatory connections between nearby cells and inhibitory connections between distant cells; this allows a bump of activity to remain stable at any position along the ring. Finally, the output connectivity of direction-selective cells is shifted along the ring, with stronger connections to nearby cells ahead in the preferred direction, whereas non-selective cells have symmetric output connectivity. This connectivity profile allows the directional cells to move the bump around the ring in response to directional input.

To determine whether the trained model resembled a canonical CBAN, we first computed, for each cell, the angle of maximum activation (*θ_max_*) during an unstructured task. We then rearranged the rows and columns of the recurrent weight matrix **W***_Rec_* according to increasing *θ_max_*. The sorted matrix revealed a pattern of local excitation (Fig. 4a). Because competition in our model is implemented through the activation function rather than explicit inhibitory neurons, the absence of long-range inhibitory connections is expected. The observed pattern is therefore reminiscent of the Mexican-hat connectivity employed in CBANs. Next, we asked whether place cells showed selectivity for the direction of motion. To address this question, we evaluated the model on CW and CCW trajectories and computed the peak activation of each cell in each direction (Fig. 4b). As in CBANs, we identified three subsets of cells: CW- and CCW-selective cells (hereafter referred to as directional cells) and non-direction-selective cells, which we refer to as central cells by analogy with the central population in CBANs. We classified cells as selective to a given direction if the maximum firing was greater that 0.01 in the preferred direction and smaller than 0.01 in the non-preferred direction; cells were classified as central if they did not belong to any of the direction-selective populations. As a result, from the 128 cells in the network, 25(20%) were classified as CW cells, 26(20%) were classified as CCW cells, and 77(60%) were classified as central. As expected, *θ_max_* matched a uniform distribution for each population (Kolmogorov-Smirnov test, D = 0.07, p = 0.70 for central cells, D = 0.16, p = 0.49 for CW cells, and D = 0.08, p = 0.98 for CCW cells, histograms shown in Figure S4).

**Figure 4:**
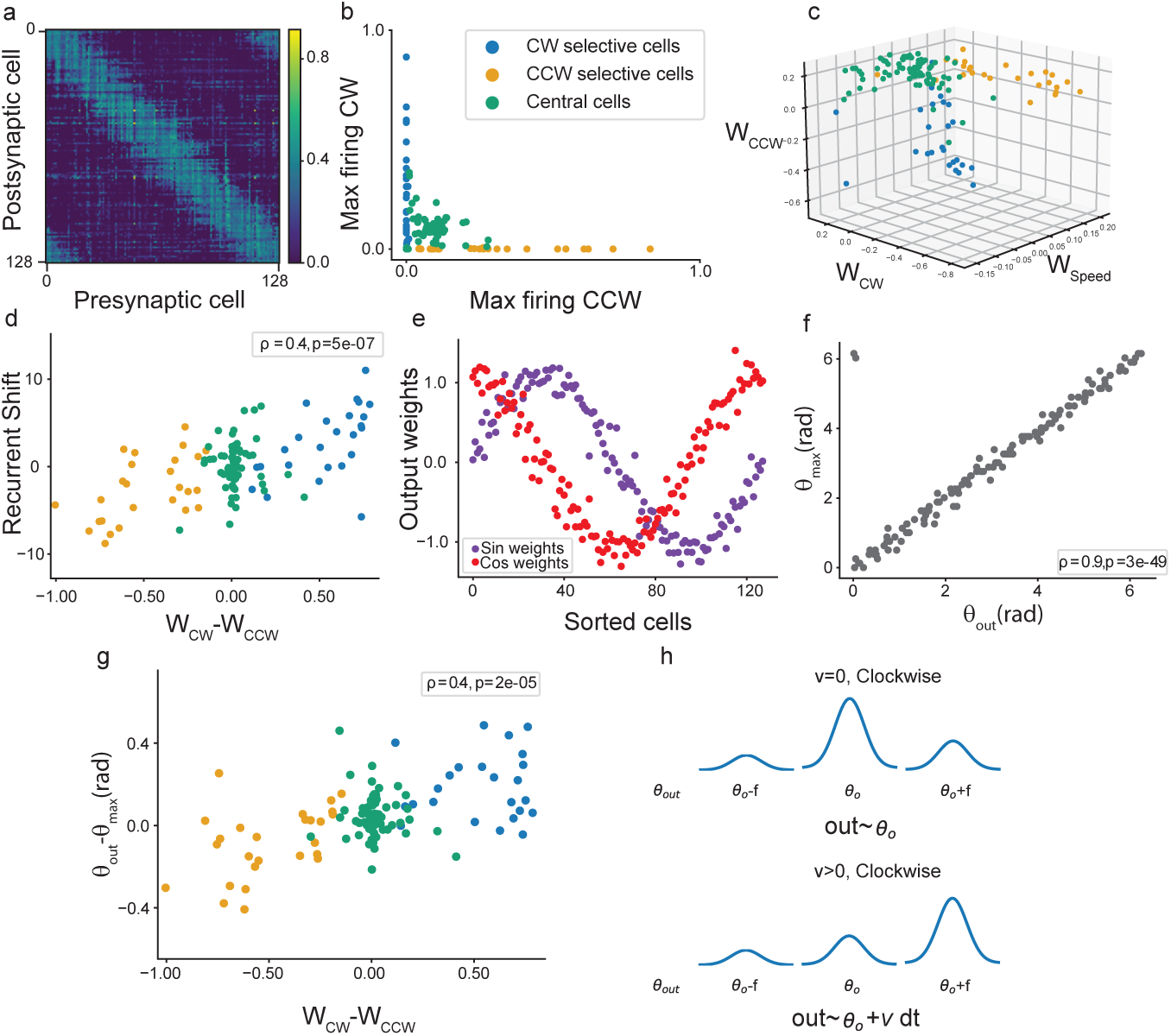
An RNN trained to solve a path-integration task resembles a CBAN. **a** Recurrent weight matrix sorted by each cell’s preferred position *θ_max_*. Columns correspond to outgoing weight vectors, and rows to incoming weight vectors. Thus, matrix entry (*i, j*) represents the weight of the connection from the neuron in column *j* to the neuron in row *i*. **b** Maximum firing for each cell in trajectories with fixed directions; cells are classified as direction-selective if they have a maximum firing *>* 0.01 in the preferred direction and < 0.01 in the non-preferred direction (20% CW, 20% CCW), and as central cells other-wise (%60). **c** Direction and speed input weights for all cells. **d** Recurrent shift, defined as the distance from the center of mass of each column to the diagonal of the matrix, plotted against the difference between CW and CCW input weights. The recurrent shift is positively correlated with the direction input weight difference, consistent with CW-selective neurons projecting preferentially toward neurons with higher *θ_max_* values and CCW-selective neurons toward lower *θ_max_* values. **e** Output weights *O*_1_, *O*_2_ corresponding to targets *sin*(*θ*), *cos*(*θ*), for all units in the network, sorted by the angle of maximum firing *θ_max_*. **f** The preferred output angle 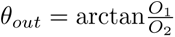 closely matches *θ_max_* for each cell. **g** Cells with stronger inputs favoring the CW or CCW direction have preferred output angles shifted ahead of or behind their preferred firing position. **h** Schematic of how the network could balance cells that are active at the same angle but have different preferred output angles, to return the correct position.

Direction selectivity can be understood from the distribution of input weights (Fig. 4c), which forms distinct clusters for each population. Although the optimized weights converged to both positive and negative values, the Softmax activation is invariant to the addition of a constant to all weights. Consequently, the signs of the weights cannot be interpreted directly in terms of excitation or inhibition. Instead, cells with above-average input weights are expected to be preferentially activated by a given input, whereas cells with below-average input weights are expected to be suppressed. Under this criterion, central cells received, on average, weaker speed input than CW and CCW cells (**W***_speed_* median = 0.05 for central cells, and 0.12 for directional cells) and are therefore preferentially suppressed by the Softmax competition as speed increases. In contrast, directional cells received similar speed input but are preferentially suppressed by the opposite direction input (**W***_CW_* median = 0.19 for CW cells, −0.34 for CCW cells,; **W***_CCW_* median = 0.15 for CCW cells, −0.46 for CW cells).

To further compare our model with classical CBANs, we asked whether directional cells also exhibited shifted recurrent connectivity. To this end, we computed the center of mass of the local excitatory connectivity profile of each cell (i.e., each column of the sorted recurrent weight matrix **W***_Rec_*, corresponding to its outgoing recurrent connections) and defined the connectivity shift as the distance between this center of mass and the matrix diagonal, which represents a connectivity profile centered on the presynaptic cell. Directional cells exhibited significant connectivity shifts (median = 4.1, Wilcoxon signed ranked test *T* = 36*, p* = 5 × 10*^−^*^4^ for CW cells, median = −2.7, Wilcoxon signed ranked test *T* = 61*, p* = 2 × 10*^−^*^3^ for CCW cells), whereas central cells displayed approximately centered connectivity profiles (median = −0.1, Wilcoxon signed ranked test *T* = 1446*, p* = 0.64). Furthermore, we defined a directional selectivity index based on the asymmetry of the direction input weights, *SI_dir_* = *W_CW_* − *W_CCW_*. We found the recurrent shifts correlated significantly with *SI_dir_* (Spearman’s *ρ* = 0.4*, p* = 5 × 10*^−^*^7^, Fig. 4d). This result is again consistent with classical CBANs and explains how activity propagates in the direction of motion: direction and speed inputs activate directional cells, whose shifted recurrent connectivity preferentially excites cells ahead of the activity bump, thereby driving its displacement.

Finally, we examined how the different cell populations are read out by the output layer. We found that the output weights closely follow sine and cosine profiles across the population (Fig. 4e). Because the network uses a sparse place-cell code, only a small subset of cells is active at any given position. Thus, the outputs (sin *θ*^^^, cos *θ*^^^) are generated by selecting and combining a small number of points along these sinusoidal profiles. This observation suggests a natural interpretation of the output weights. Since cells contribute to both the sine and cosine outputs, each cell can be assigned a preferred output angle, 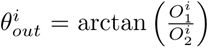 where 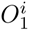 and 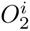 are its weights onto the sine and cosine outputs, respectively. We found that *θ_out_* was strongly correlated with the position at which the cell was maximally active, *θ_max_* (Fig. 4f; Spearman’s *ρ* = 0.86*, p* = 3 × 10*^−^*^49^). Thus, the preferred output angle of a cell is largely determined by its preferred position. Nevertheless, the relationship was not exact, and many cells exhibited a noticeable difference Δ*θ* = *θ_out_* − *θ_max_*. To determine whether these differences were related to directional selectivity, we studied the relationship between the selectivity index *SI_dir_* and Δ*θ*, and found a positive correlation (Fig. 3g; Spearman’s *ρ* = 0.4*, p* = 2 × 10*^−^*^5^). Thus, cells with similar *θ_max_* can be coactive while contributing to slightly different output angles. Because the relative activity of these cells is modulated by the velocity inputs, changes in movement direction can re-balance their contributions to the output and thereby produce instantaneous shifts in the represented angle. This is schematized in Fig. 4h, for a case with only one cell of each subpopulation. In this example, all three cells are selective to an angle *θ*_0_; the central cell has an output angle *θ*_0_, while directional cells have a shift ±*f* between *θ_max_* and *θ_out_*. In the absence of velocity input, their combined contribution yields an output angle *θ*_0_. However, when the speed input suppresses the central cells, the balance is altered and the represented angle shifts clockwise. The same mechanism extends naturally to broader activity bumps, where the represented angle emerges from the combined contributions of all active cells, each with its corresponding output-angle shift.

Taken together, these results show that optimizing an RNN for path integration naturally gives rise to a CBAN-like mechanism. The learned network recapitulates the key structural and dynamical features of classical CBANs, and the output layer exploits this organization to decode position. We next investigate whether specific features of this learned architecture can explain the concave speed response.

### 2.5 Concave speed response emerges from a central-cell brake mechanism

Because speed inputs preferentially suppressed central cells, and changes in the balance between central and directional populations were associated with bump displacement, we hypothesized that the concave speed response observed in Fig. 3f arises from the progressive silencing of the central population as speed increases. To test this hypothesis, we measured the mean activity of each cell population during CW trajectories in which speed was held constant within each episode and systematically increased across episodes. Central-cell activity decreased with increasing speed, whereas CW-cell activity increased (Fig. 5a, fully opaque traces). Importantly, the increase in CW-cell activity was itself concave. These observations suggest the following mechanism. Because central cells have unshifted recurrent connectivity, they tend to stabilize the activity bump rather than move it. As speed increases, the speed input progressively suppresses the central population, reducing its stabilizing influence and allowing directional cells to drive bump motion more effectively. However, this suppressive effect is inherently bounded: once most central cells have been silenced, further increases in speed produce progressively smaller changes in the balance between the two populations. Consequently, the increase in directional-cell activity—and therefore attractor speed—becomes progressively smaller, giving rise to the observed concave speed response. We refer to this as the central-cell brake mechanism, because the central population acts as a brake on the movement of the activity bump.

**Figure 5:**
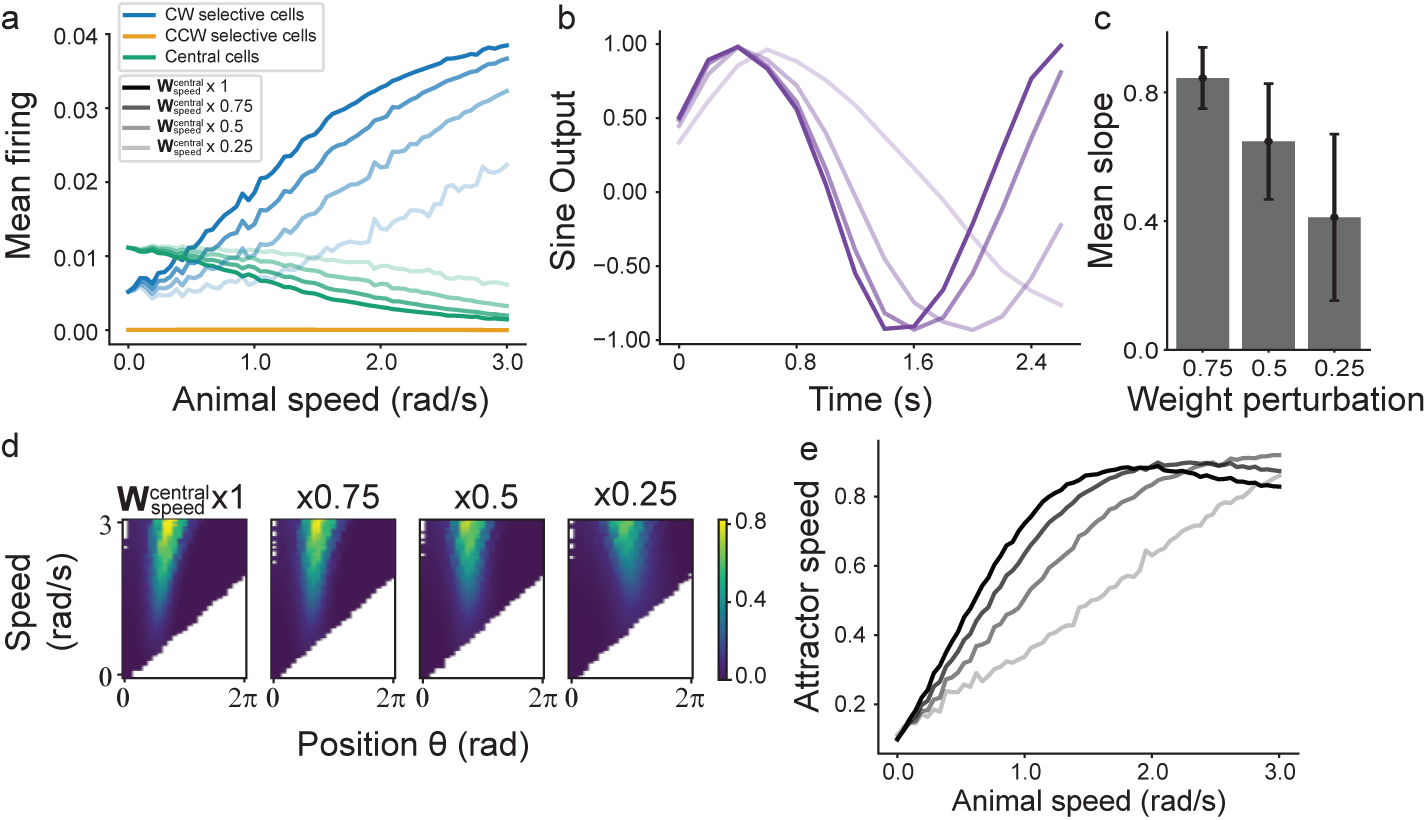
Perturbation analysis reveals a central-cell brake mechanism underlying the concave speed response. **a** Mean firing of each of the three cell populations, for the original model (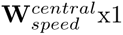, fully opaque lines) and after progressive reduction of speed inhibition to central cells, by translating and dividing **W**_speed_ (see methods). **b** Examples of the model’s sine output for CW trajectories at v = 3 rad/s, shown for the four perturbation levels. Increasing perturbation stretches the output in time. **c** Slope of linear fits between the angle estimated by the unperturbed model, and each manipulated model (mean±std). Pearson’s r > 0.98 in all trials. **d** Example speed-ordered spatial rate maps for the original model and progressively perturbed models, showing a progressive reduction in place-field shifts for the same cell. e Attractor speed versus animal speed for the original model and progressively perturbed models, showing a progressive loss of the concave speed response and an increasingly linear relationship.

To test the proposed brake mechanism, we progressively weakened the connections from the speed input to the central-cell population by multiplying its corresponding entries of **W***_speed_* by 1, 0.7, 0.5, and 0.25 (see Methods). We then repeated the fixed-speed experiments described above, in which CW trajectories were evaluated at constant speed within each episode while speed was systematically increased across episodes. As expected, increasingly stronger perturbations produced progressively slower silencing of the central cells and a slower, less concave increase in CW-cell activity (Fig. 5a). Importantly, these perturbations did not disrupt path integration. Instead, the network generated essentially the same trajectories, but at a reduced speed (Fig. 5b). To quantify this effect, we compared the decoded angles from perturbed and unperturbed networks. For each perturbation, the decoded angles remained highly correlated with those of the unperturbed network (Pearson’s *r >* 0.98), whereas linear regression revealed a decrement in slope with increasing perturbation strength, reflecting the progressive stretching of the estimated trajectories. (Fig. 5c). Furthermore, firing-field shifts became smaller (shown for one example cell in Fig. 5d), and the relationship between attractor speed and animal speed became increasingly linear (Fig. 5e). Together, these results provide causal support within the model for the proposed central-cell brake mechanism and strengthen the link between central-cell silencing, the concave speed response, and the emergence of firing-field shifts.

### 2.6 The connectivity motifs identified in the trained network are sufficient to reproduce the concave speed response and firing-field shifts

To test whether the recurrent and input connectivity motifs identified in the previous sections are sufficient to reproduce the concave speed response and firing-field shifts, we constructed a simplified recurrent network incorporating only these features. Each cell was assigned a Gaussian output connectivity profile of width *s*, centered on the cell itself for central cells or shifted by an amount *f* for CW and CCW cells. The recurrent weight matrix **W***_Rec_* was then constructed by assigning these connectivity profiles as its columns, with cells ordered as repeating triplets of CCW, central, and CW neurons. Input weights were chosen according to the average connectivity profiles measured in the trained network (Fig. 6b). Specifically, the speed input targeted only central cells through a tunable weight **W***_speed_*, whereas directional inputs inhibited the opposite directional population. The Gaussian connectivity parameters (*A*_0_*, A_C_, s, f*) and the Softmax inverse temperature *β* were also tuned. Each episode was initialized with a localized activity bump consisting of unit activity in the first central neuron and zero activity elsewhere.

**Figure 6:**
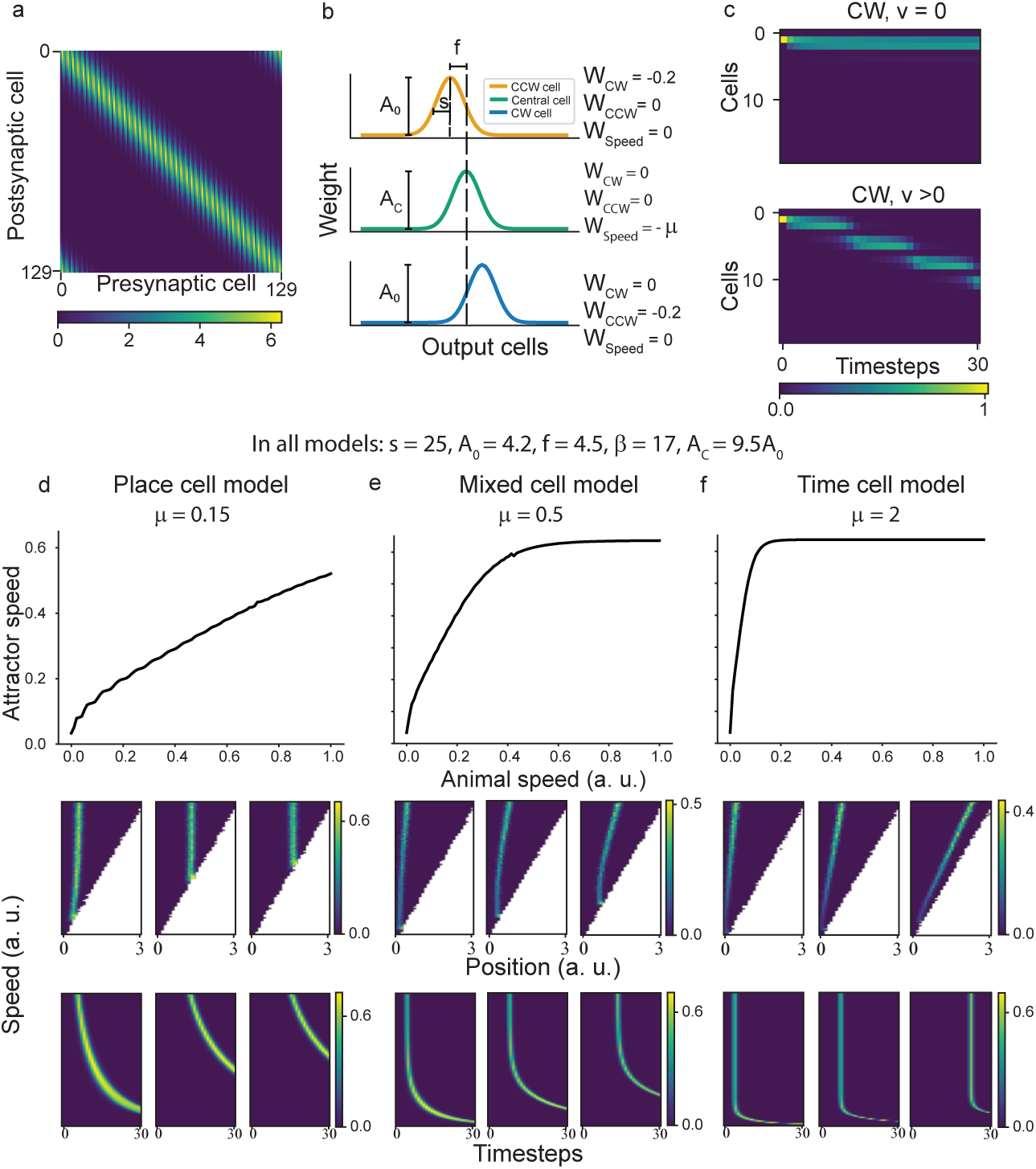
A simplified RNN reproduces the concave speed response and firing-field shifts. **a.** Connectivity of the simplified model. Columns correspond to outgoing weight vectors, and rows to incoming weight vectors. The matrix alternates CCW, central, and CW cells. **b** Output recurrent weights for directional and central cells are defined as shifted gaussians with different amplitudes *A*_0_ and *A_C_* respectively. Directional cells are inhibited by the direction input that is opposite to its preferred direction, and central cells are inhibited by speed **c** Activity heat maps for a simplified model with *µ* = 0.5 and CW input. Top: With zero speed, the activity bump remains nearly stationary. Bottom: With positive speed the activity bump moves along the CW direction. **d** With *µ* = 0.15 the model becomes a place-cell model. It exhibits place fields and an approximately linear relationship between attractor speed and animal speed. **e** With *µ* = 0.5 the model becomes a mixed-selectivity cell model. It exhibits shifting place and time fields as seen by Chen et al., and a concave attractor speed response. **f** With *µ* = 0.2 the model becomes a time-cell model. It exhibits time fields and an approximately constant relationship between attractor speed and animal speed this model, elapsed time was not represented by a separate population of pure time cells, but by cells with joint space-time fields. This resembles experimental observations in which hippocampal time cells can also be modulated by position [11], although this particular model did not produce pure place or pure time cells.

Across a range of parameter values, the simplified network successfully reproduced controlled bump motion in both speed and direction, and was able to path-integrate (Fig. 4c). The parameter **W***_speed_* controlled the qualitative coding regime of the network. Small absolute values of **W***_speed_* produced an approximately linear attractor-speed response, together with pure place fields and shifting time fields (Fig. 6d). Intermediate values generated a concave, saturating speed response and firing-field shifts in both the spatial and temporal reference frames (Fig. 6e). Finally, large absolute values caused attractor speed to saturate at low animal speeds, giving rise to pure time fields and shifting place fields (Fig. 6f). This behavior is consistent with both the perturbation experiments in the trained network, and the effects of varying *η* in the line-attractor model.

Taken together, these results show that the recurrent and input connectivity motifs identified in the trained network are sufficient to reproduce the family of concave speed responses considered in the line-attractor model and the corresponding firing-field shifts. Thus, the same computational principle emerges consistently across the three levels of modeling considered so far.

### 2.7 An RNN trained to encode space and time develops genuine joint space-time fields

The previous sections showed that the place and time-field shifts reported by Chen et al. can arise in models trained to encode space alone, provided that space and time are correlated at the behavioral level. We next asked what a genuine space-time representation would look like in a recurrent network explicitly trained to encode both variables. To this end, we trained an RNN to simultaneously solve two independent tasks: the one-dimensional path-integration task described above, and a timing task in which the network received a one-hot cue at random times and had to report the elapsed time since the last cue (see Methods). Figure 7a shows an example episode with two cue presentations and the corresponding model output. Importantly, the spatial and temporal tasks were independent by design. Thus, the network could in principle represent position and elapsed time separately, and any coupling between the two representations reflects a strategy developed during training rather than a constraint imposed by the task.

**Figure 7:**
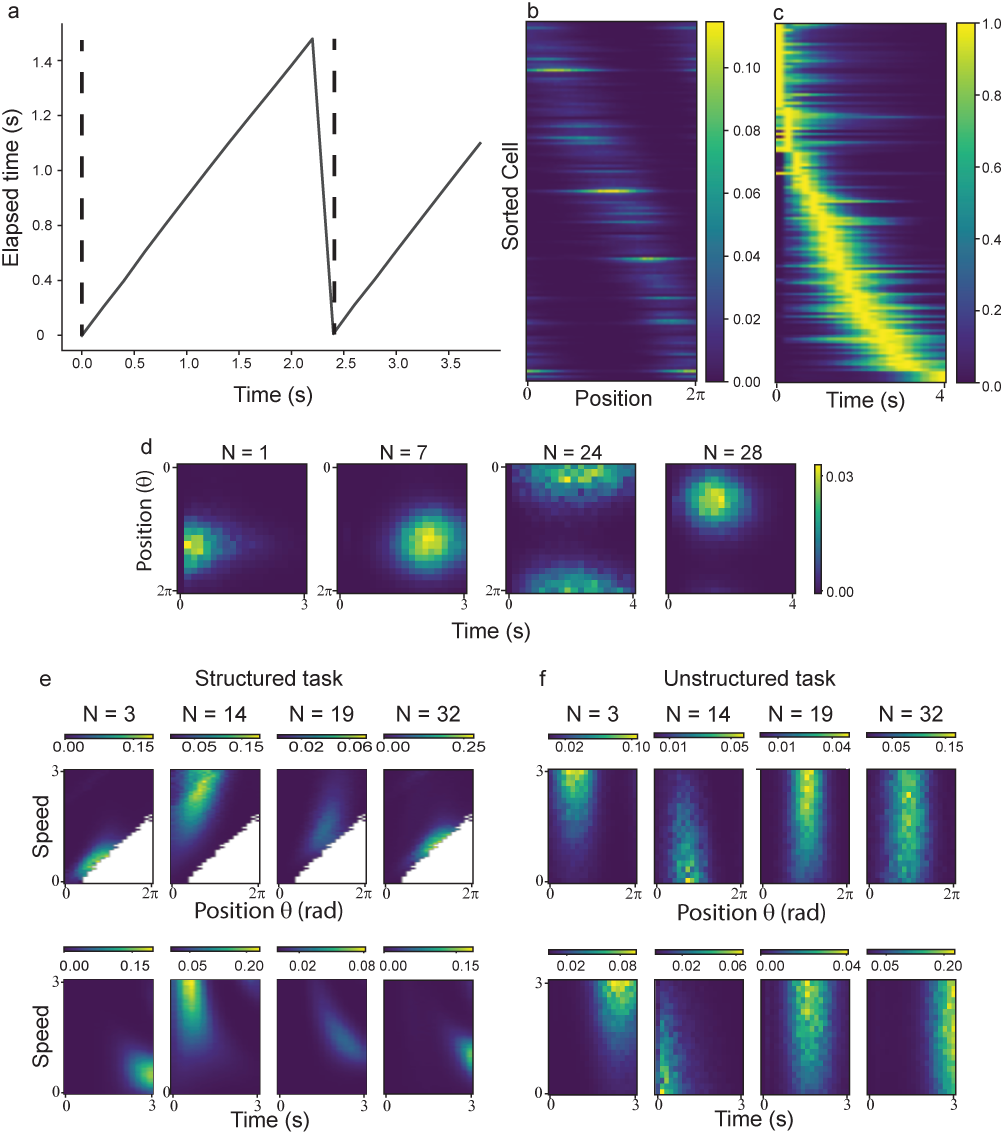
An RNN jointly trained for spatial and temporal encoding develops mixed-selectivity representations. **a** Trial example of the timing task with two cues presented, and time integrated from the previous stimulus. **b** Sorting rate maps by the position of maximum firing reveals place cells that tile the entire track. **c** Normalized average activities over trials at each time step, revealing time cells that tile the entire length of the task. **d.** Position-time rate maps for some example cells show that most cells encode both time and space. **e** Top: Speed-sorted rate maps show place fields shifting forward but with an amplitude modulation, in trajectories starting from the same initial position and with the same direction. Bottom: Speed-sorted trial average plots show encoding of time with a backward shift and an amplitude modulation. **f.** Same as (e) but averaging trajectories starting from different initial positions.

After training, we analyzed how cells in the recurrent layer encoded space and time. Spatial rate maps computed on random trajectories and sorted by the position of maximum firing revealed that cells tiled the entire track (Fig. 7b). Compared with the network trained only for spatial encoding, these fields were broader and more variable in amplitude, possibly reflecting the additional demand of encoding time simultaneously. We then evaluated temporal tuning by averaging activity over episodes with a single cue presentation at *t* = 0. This analysis revealed that cells also tiled the full temporal interval (Fig.7c). Finally, to characterize joint coding, we computed two-dimensional space-time firing maps. As illustrated for four example cells in Fig. 7d, cells fired preferentially at specific combinations of position and elapsed time, and their field centers covered both dimensions (all two-dimensional maps are shown in Fig. S10). Thus, in

We then asked whether this model would behave differently from the space-only models when evaluated using the structured and unstructured protocols described above. In structured trajectories with fixed initial position and a cue presented at *t* = 0, both place and time fields shifted with running speed (Fig. 7e). However, when initial position, speed and direction were randomized, temporal selectivity remained evident while field shifts disappeared (Fig. 7f). Thus, unlike the apparent time fields observed in the space-only models, temporal tuning in this network persisted when the correlation between space and time was broken. This provides a concrete example of genuine elapsed-time coding and how the unstructured task distinguished it from the apparent temporal tuning induced in the structured task.

We next examined whether the spatial component of this joint code relied on mechanisms similar to those found in the network trained only for path integration. Overall, the same signatures were present, although they were noisier (Fig. S9). Sorting the recurrent weight matrix by the position of maximum firing revealed a local excitation pattern. Fixed-direction trajectories revealed CW-selective, CCW-selective, and central cells. The output weights again followed sine and cosine profiles, although the relationship was more variable than in the space-only model. The preferred output angle, *θ_out_*, was strongly correlated with the position of maximum firing, *θ_max_*, and their differences correlated with selectivity. These results indicate that the network solved the spatial task using a CBAN-like mechanism similar to that characterized above. One feature was less pronounced, however: unlike the space-only model, the center of mass of each cell’s recurrent output connectivity did not correlate significantly with direction selectivity (Spearman’s *ρ* = −0.1, *p* = 0.17). This difference is consistent with the additional constraint imposed by the joint task, in which the recurrent connectivity must simultaneously support path integration and the autonomous bump motion required for temporal coding. Importantly, the remaining signatures of the CBAN-like mechanism were preserved.

We then analyzed the mechanism supporting elapsed-time encoding. Sorting the recurrent matrix by the time of maximum firing revealed a local excitation pattern among time cells, except for cells active at the first few time steps (Fig. 8a). In addition, the center of mass of the recurrent output connectivity was significantly shifted toward positive values (Median shift = 4.3, Wilcoxon signed-rank test, *T* = 2014*, p* = 5 × 10*^−^*^7^), consistent with a mechanism that propagates activity through a temporal sequence (Fig. 8b). Cue-related inputs provided a complementary mechanism for resetting this sequence. Cells representing early times received positive cue input, whereas cells representing later times received negative cue input (Fig. 8c). This organization allows the cue to restart the temporal sequence independently of which cells were active when the cue arrived. Finally, we asked how the output layer converted time-cell activity into an estimate of elapsed time. The output weight of each time cell was strongly correlated with the time at which that cell fired maximally, and in many cases closely matched that time (Spearman’s *ρ* = 0.96*, p* ≪ 0.001, Fig. 8d). Although multiple cells encoded similar elapsed times, they were active at different positions, so only a small subset contributed to the output at any given moment. Together, these analyses suggest a simple temporal computation: cue inputs reset the sequence, shifted recurrent connectivity propagates activity through time, and the output layer maps the active cells onto the corresponding elapsed time.

**Figure 8:**
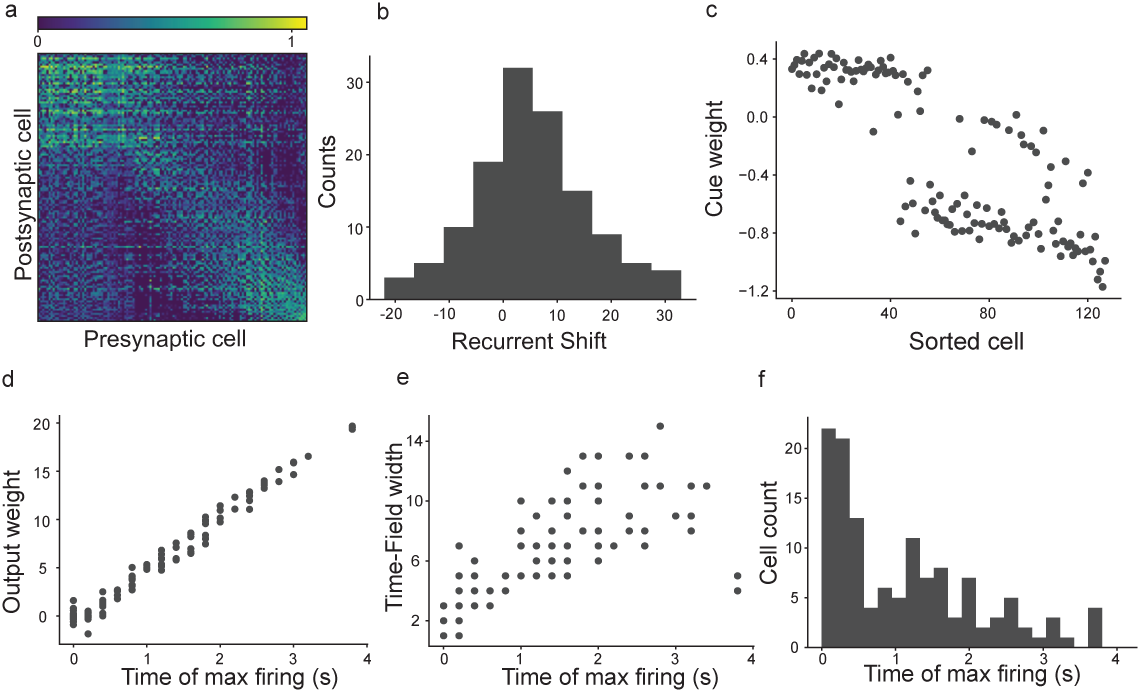
Connectivity underlying joint spatial and temporal representations. **a** Sorting weights according to the time of the maximum firing for each cells reveals a local excitation pattern for cells that fire at the beginning of the sequence. **b** Recurrent shift, defined as the distance from the center of mass of each column in **W***_Rec_* to the diagonal of the matrix, are slightly shifted forward in the sequence. **c** Cue input excites cells that fire in the first part of the sequence, and increasingly inhibits cells that fire at latter times. **d** Output weights select the time when each time cell fires maximally. **e-f** Time cells in the model show evidence of Weber’s Law, with a positive correlation between time of firing and time field width (e), and more cells representing earlier than late times (f).

Having established that the model genuinely represented elapsed time, we asked whether its temporal code exhibited properties observed in biological time cells. Time-cell populations have been proposed to use a logarithmically compressed representation of time, producing two characteristic signatures: more cells fire early in the interval than late, and later time fields are wider [33]. We found both signatures in the model. Defining time-field width as the number of time steps above half-maximal activity, we found a positive correlation between peak time and field width (Spearman’s *ρ* = 0.83*, p* ≪ 0.001, Fig. 8e). In addition, more cells represented early elapsed times than late elapsed times (Fig. 8f).

Thus, although the model was trained only to estimate elapsed time, its temporal representation exhibited Weber-like properties similar to those reported experimentally. Taken together, these results provide a contrasting case to the space-only models analyzed above. In those models, temporal tuning was apparent and disappeared when the behavioral correlation between space and time was removed. Here, by contrast, a network trained to encode both variables developed joint space-time fields and retained temporal selectivity under decorrelated trajectories. This model therefore illustrates one way in which genuine elapsed-time coding can coexist with spatial coding while still producing speed-dependent field shifts reminiscent of those reported by Chen et al.

## 3 Discussion

Space and time encoding is essential for navigation, action organization, and episodic memory, and hippocampal populations are thought to contribute to both forms of representation. Recent experimental work suggested that CA1 neurons jointly represent space and time, and that their place and time fields shift in a coherent, speed-dependent manner [18]. Here we developed a series of models that support an alternative possibility: firing-field shifts can arise when the behavioral task correlates space and time, and when the internal spatial representation is supported by a continuous attractor whose speed does not scale proportionally with the animal’s speed. We reproduced the place- and time-field shifts reported by Chen et al. using a simple line-attractor model, and the same behavior emerged in recurrent neural networks optimized to encode space, despite lacking an explicit elapsed-time output. These results do not preclude the existence of neurons with genuine mixed selectivity for space and time in CA1. Rather, they show that field shifts alone cannot be taken as evidence for an explicit space-time interaction, or for competition between spatial and temporal coding. In this sense, the models introduced here provide a null hypothesis for interpreting joint space-time responses.

When space and time are correlated by the structure of the task, a purely spatial representation can produce apparent temporal fields and speed-dependent shifts. Conversely, when initial position and movement direction are randomized, this behavioral correlation is removed and the apparent temporal tuning generated by the space-only models disappears. This provides a simple experimental criterion: if temporal tuning reflects spatial dynamics under constrained behavior, it should be strongly reduced when the relationship between position and elapsed time is decorrelated. If, instead, neurons genuinely encode elapsed time, temporal selectivity should persist under such conditions. Our modeling work therefore highlights the importance of carefully disentangling behavioral variables when interpreting mixed selectivity, especially in tasks in which position, elapsed time, speed, sensory input, and task phase are naturally correlated [15–17].

The mechanism we identified is closely related to continuous bump attractor models of spatial representation and path integration [30–32]. In classical bump attractor models, a population of neurons represents the encoded variable, such as position or head direction, through a localized activity bump. Short-range excitation and long-range inhibition stabilize the bump, whereas velocity-related inputs introduce asymmetries that move it through the representational space. In many hand-designed attractor models for path integration, bump velocity is constructed to scale approximately linearly with velocity input [30,34–37]. This proportionality often follows from the symmetry and connectivity assumptions that make the models analytically tractable, and it is required for accurate path integration over a broad range of speeds. In the present setting, however, a perfectly linear gain cannot explain the observed field shifts. Instead, the relevant regime is one in which the velocity of the internal spatial representation becomes progressively less sensitive to the velocity of the animal.

The RNNs optimized for spatial encoding recapitulated several organizational principles of continuous bump attractors, but did so in a more distributed form. Neurons developed location-selective responses that tiled the track and formed an activity bump, and the recurrent connectivity showed strong local structure. At the same time, the trained networks did not rely on sharply separated functional populations or a rigid one-to-one mapping between direction-selective inputs and position-encoding neurons. Instead, they exhibited a continuum of direction selectivities, ranging from strongly CW-selective to strongly CCW-selective neurons, with many weakly selective cells in between. Direction selectivity was aligned with the displacement of each neuron’s recurrent output connectivity, with CW- and CCW-selective neurons driving the bump in opposite directions. Thus, optimization recovered a CBAN-like organization while relaxing some of the strong architectural constraints typically imposed in hand-designed models [30,31]. This distributed organization also influenced how directional information was represented. Direction inputs rapidly rebalanced activity within the bump, increasing the contribution of direction-selective neurons before the bump center had fully shifted. This effect helps explain how the trained network could update its position estimate during rapid changes in movement direction, and illustrates that motion variables can be represented through graded changes in population activity rather than through sharply defined direction-specific modules.

The central mechanism underlying the speed-dependent field shifts, however, was a brake mechanism implemented by a weakly direction-selective, stabilizing population. In the trained networks, these cells had nearly symmetric recurrent connectivity and therefore contributed little to bump motion. Instead, they tended to stabilize the activity bump. Because the model used a Softmax non-linearity, the absolute sign of individual input weights cannot be interpreted directly as biological excitation or inhibition. The relevant observation is the relative ordering of speed weights: increasing speed reduced the influence of the weakly selective population relative to more direction-selective neurons. As speed increased, the stabilizing population progressively released its brake on bump motion, allowing neurons with shifted recurrent connectivity to exert a greater influence on the network state. This redistribution of influence provides a circuit-level implementation of the concave relationship between animal speed and attractor speed assumed in the line-attractor model.

The brake mechanism generates several experimentally testable predictions. First, if a similar attractor-like organization is present in hippocampal or entorhinal circuits, weakly direction-selective neurons should show a stronger reduction in activity with increasing speed than neurons with stronger directional tuning. Second, direction selectivity, recurrent connectivity shift, and speed sensitivity should be related: cells with stronger directional preference should tend to have more shifted effective connectivity and a different speed dependence than weakly selective cells. Third, population-level estimates of internal bump velocity should scale sublinearly with running speed in conditions that produce field shifts. Finally, manipulations that prevent the speed-dependent reduction in the influence of the weakly selective central-like population should reduce the concavity of the attractor-speed response and diminish the corresponding place and- time-field shifts. This last prediction differs from a simple suppression experiment: the key manipulation is not to silence the stabilizing population per se, but to block or reduce the speed-dependent change in its influence.

Several known properties of hippocampal-entorhinal circuits make this mechanism biologically plausible. Speed signals are present in the hippocampal formation in several forms, including theta modulation [38], pure speed cells [39], and conjunctive cells that encode speed together with direction, position, or other variables [7]. Some speed responses are approximately linear [39], whereas others are sublinear or saturating [40]. Such sublinear signals are consistent with the concave speed response analyzed in our models, but they do not by themselves explain how this response could arise at the circuit level. The trained RNN suggests one possible implementation: speed may alter the relative influence of stabilizing and motion-driving populations. Medial entorhinal cortex contains speed and direction signals and projects to hippocampal circuits [7, 41], and speed-modulated inhibitory projections from entorhinal cortex to hippocampus have been reported [41,42]. Although our model does not allow the sign of individual speed weights to be interpreted directly, the preferential reduction in the influence of weakly selective cells is qualitatively consistent with the possibility that speed-dependent inhibitory pathways regulate the contribution of specific hippocampal or entorhinal populations.

After establishing how apparent temporal tuning can arise from spatial dynamics, we trained a second class of networks to jointly encode position and elapsed time. This model served as a contrasting case: unlike the space-only networks, it contained a genuine temporal output. The resulting representation was mixed. Neurons fired preferentially at specific combinations of position and elapsed time, resembling experimental observations in which hippocampal time cells are modulated by spatial location or behavioral context [11, 15]. The model also exhibited signatures associated with Weber-like temporal coding: later time fields were broader, and more cells represented early than late elapsed times [33]. These results show that genuine elapsed-time coding can emerge in the same general modeling framework, but with a different diagnostic signature. When tested under a protocol resembling that of Chen et al., the joint space-time network exhibited speed-dependent field shifts. However, when position and elapsed time were decorrelated, the shifts disappeared while temporal selectivity remained. Thus, decorrelation distinguishes apparent temporal tuning induced by spatial dynamics from genuine elapsed-time coding.

The present work has some limitations. First, the proposed mechanism should be regarded as one possible circuit implementation capable of explaining the observations of Chen et al., rather than definitive evidence that hippocampal circuits operate in this way. Other recurrent architectures may reproduce the same phenomena through different circuit organizations. Second, the Softmax competition used in the recurrent dynamics prevents a direct biological interpretation of the sign of individual synaptic weights. Consequently, the model predicts relative changes in the influence of different neuronal populations with speed, rather than excitatory or inhibitory interactions at the level of individual synapses. Third, the recurrent network trained to jointly encode space and time produced predominantly mixed-selectivity neurons, and therefore does not capture the full diversity of hippocampal responses, including distinct populations of pure place and pure time cells [18]. Finally, the models were developed for one-dimensional navigation, and whether the same mechanisms account for mixed space-time representations in more complex behavioral settings remains an open question.

Taken together, our results provide an alternative interpretation of the observations reported by Chen et al. Rather than requiring an explicit representation of elapsed time, the observed field shifts and apparent space-time competition are consistent with a nonlinear, concave relationship between animal speed and the velocity of an internal spatial representation of position. The trained recurrent networks further suggest a concrete circuit implementation of this phenomenon through a central-cell brake mechanism, in which a weakly direction-selective population stabilizes the attractor at low speeds and progressively releases this constraint as speed increases. Beyond explaining the existing observations, this mechanism extends classical bump attractor models with a new functional role for the stabilizing population [30–32] and provides experimentally testable predictions linking direction selectivity, speed sensitivity, recurrent connectivity, and the neural dynamics underlying path integration.

## 4 Methods

### 4.1 Pure place cells or time cells cannot explain field shifts

To simulate both place and time cells, we defined their activity as a gaussian function of position or elapsed time, respectively. For the structured task, we initiated 100 trajectories at position 0, each one with a different speed, ranging from 0 to 1, always moving to the right.

For the unstructured task, we randomly chose initial positions between 0 and *L* and sampled speed from a standard-normal at each time-step, with the sign defining direction of movement. Time was still measured since the start of the trial.

### 4.2 Shifts in place and time fields explained with a continuous line attractor model

We considered a mouse running in a linear track whose position *p* is measured from the starting point, located at the origin of the track (*p*(*t* = 0) = 0). The position at time *t* is the solution to the integral 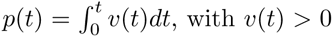 the speed of the animal. We will assume that, during a single run, the mouse’s speed is constant, hence we have that *p*(*t*) = *vt*. The neural representation of the mouse’s position can be modeled in terms of a continuous line attractor. This kind of attractor can be approximated by a neural network with a bump attractor dynamics. For simplicity, here we will consider a continuous linear attractor defined by 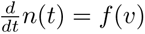. The variable *n*(*t*) encodes the active neuron at time *t*. Its derivative is a function of the animal’s speed, linking changes in position with changes in the neural representation of this position. We will further assume that *n*(*t* = 0) = 0, and that *f* (*v*) is a positive, non-decreasing and concave function. Since *v* is constant, 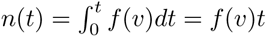Therefore, a given neuron n will be activated at time 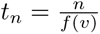, and the animal’s position during activation will be 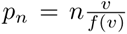. It is easy to see how speed affects the timing of activation. As *v* increases, *t_n_* decreases, replicating the shift towards earlier times seen in Chen et al., where laps are ordered in increasing speed. To see the effect of speed on the animal’s position at which *n* is activated, we differentiate *p_n_* with respect to 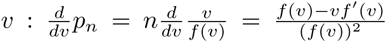, which is positive if *f* (*v*) − *vf^′^*(*v*) *>* 0. By the mean value theorem, there exist *c* ∈ (0*, v*) such that 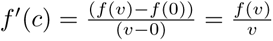 Since *f^′′^*(*v*) *<* 0 (because *f* (*v*) is concave), then 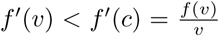. Therefore, *vf^′^*(*v*) *< f* (*v*), proving that *p_n_* increases with *v*, replicating Chen’s results where place field shifted towards the end of the track when the animal run faster (decreasing range duration). If we further assume that *f* (*v*) is bounded, then *t_n_* tends to a linear function of *n* as speed increases, hence neurons tend to be perfect time cells in this scenario.

### 4.3 Competition between space and time encoding

In the setting of Chen et al. the beginning of the lap is the origin from which both, position and time, are measured. Within the track, animals can only move forward from this frame of reference. This makes space and time utterly correlated. If we say that a neuron is both a place cell and a time cell, it means that it activates in a particular place and time from the lap start. This can only happen at a specific speed (assuming a constant speed throughout the lap). Conversely, if a neuron activates at a range of velocities (as in Chen et al. work), a trade-off between being pure place cell and pure time cell must be taking place. To see this trade-off in the context of a line attractor, let’s consider the function *f*described before (positive, non-decreasing and concave) and expand it by adding a parameter *η* which controls how much the function departs from the identity function. Let’s suppose it is also bounded. Specifically, we will assume that *f* (*η, v*) → *v* if *η* → 0 and *f* (*η, v*) → *c* if *η* → +∞, with *c* a positive constant. This in turn implies that 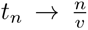 and *p_η_* → *η* when *η* → 0, and that 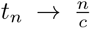 and 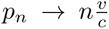 When η → +∞. In words, when the attractor speed follows the animal’s speed closer (low *η*), the position of activation becomes more independent of speed, and the time of activation tends to be a linear function of the reciprocal of speed (the neuron becomes more like a place cell). Conversely, as the attractor speed saturates with increasing animal speed (high *η*), the time of activation becomes more independent of speed, and the position of activation tends to be a linear function of speed (the neuron becomes more like a time cell).

### 4.4 Continuous line attractor implementation

For plots in Fig. 2 we computed *p_n_* and *t_n_*, taking 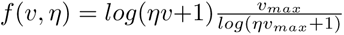, and setting *n* and a range of values for *η* to obtain curves that resemble some of the examples shown in Chen et al. The factor multiplying the logarithm is a constant that normalizes *f* output such that *f* (*v_max_, η*) = *v_max_*. This choice facilitates analysis, but has no critical role in the results obtained.

### 4.5 Recurrent neural networks trained to encode space shows shifting fields and space-time trade-offs

#### 4.5.1 Spatial training task

For each trial in the spatial task, the initial position was sampled uniformly along the track. At each time step, movement direction was sampled with equal probability between clockwise and counterclockwise motion. Speed was sampled from a normal distribution with probability (0.9) and was set to zero with probability 0.1, encouraging the network to maintain a stable position estimate while the animal remained stationary. At each time, the angle is updated according to

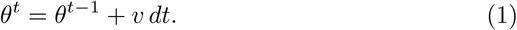

The model received the absolute value of speed, together with a one-hot input indicating if the direction of movement was clockwise or counter-clockwise. To provide landmark information, the circular track was divided into 30 angular bins. Each bin was associated with a 3x3 RGB visual pattern extending from the current bin in the direction of motion. During training, visual inputs were presented as a flattened array of 27 elements at the first timestep of every trial, providing information about the initial position, and at four additional timesteps selected uniformly at random during the trial.

At every time step, the target output *y^t^* was defined as:

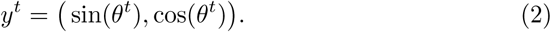

The simulation hyperparameters are summarized in Table 4.5.2.

#### 4.5.2 Network Architecture

We used a vanilla RNN [43] composed of *N* neurons, which evolved following the equation:

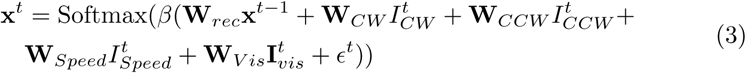

where **x***^t^*∈ R*^N×^*^1^ is the column vector of activations at time *t,* **W***_Rec_*∈ R*^N×N^* is the recurrent matrix, **W***_CW_,* **W***_CCW_ ,* **W***_Speed_*∈ R*^N×^*^1^ and **W***_vis_*∈R*^N×^*^27^ are the input weights, 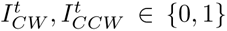 are the one-hot direction inputs, 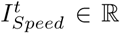 is a scalar speed input, 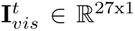 is the flattened RGB 3 × 3 visual input, *β* is an inverse temperature parameter which we used to control sparsity in the activations, and *ɛ* is normally-distributed random variable only used in training. The output of the model **y**^*^t^* is read by a linear decoder: **Ox***^t^* = **y**^*^t^*.

The network was trained by minimizing:

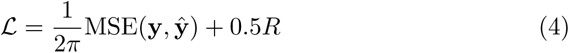

where the first term is the scaled Mean Squared Error between the output of the model and the simulation, and *R* is a custom regularizer, defined as

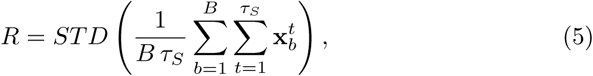

where the standard deviation is taken across cells, *b* is the mini-batch index, *B* is the mini-batch size, and *τ_S_* the number of simulated time steps in the spatial task. In other words, the regularizer promotes similar mean firing rates across cells, discouraging solutions in which a large fraction of cells remain nearly silent. Parameters **W***_Rec_,* **W***_CW_,* **W***_CCW_,* **W***_Speed_* and **W***_vis_* were optimized using backpropagation through time with ADAM as optimizer. The model was implemented in Pytorch 2.6.0.

### 4.6 Trajectory generation

To characterize the model under conditions analogous to the fixed-speed experiments of Chen et al., we generated a structured task in which every trial started at *θ* = 0, proceeded clockwise, and was performed at a constant speed. Across trials, speed was varied linearly from 0 to 3 rad/s. These trajectories were used to compute the spatial and temporal rate maps in the structured task. The visual input was only presented at *t* = 0 and, to decouple dynamics from the transitory effect produced by the visual input, each trajectory began with five time steps at *v* = 0, which were excluded from all analyses.

To decorrelate position and elapsed time, we generated an unstructured task in which the initial position was sampled uniformly along the track, while movement direction and speed were sampled independently at each time step. Direction was sampled uniformly between clockwise and counterclockwise direction, and speed was sampled uniformly from the range (0, 3) rad/s. These trajectories were used to construct the rate maps in the unstructured task, and to compute each cell’s preferred position, *θ_max_*.

Attractor speed was estimated using trajectories generated as in the structured task, except that the initial position was sampled uniformly along the track rather than fixed at *θ* = 0.

### 4.7 Speed-ordered spatial and time rate maps

The rate maps shown in Fig. 3c were computed using the unstructured task described above. For each cell, activity was averaged within spatial bins and the resulting rate maps were sorted according to the position of maximum mean firing.

The spatial rate maps in the top panels of Fig. 3d were generated using the structured task by averaging the activity of each cell across 32 trajectories with dithered initial positions to avoid empty spatial bins. All trajectories had fixed speed and clockwise direction, with different rows corresponding to speeds varying linearly from 0 to 3 rad/s. The bottom panels shows the temporal rate-maps of the same cells averaged across the same set of trajectories. Figure S2 shows the corresponding plots for all cells in the network.

The panels in Fig. 3e were generated using the unstructured task. The top panels show position vs. speed rate maps, whereas the bottom panels show elapsed time vs. speed rate maps.

Figures 4b and 5a were generated using trajectories with fixed speed and direction, but with speed varying linearly from 0 to 3 rad/s. Cells were classified as selective for a given direction if their maximum firing exceeded0.01 in the preferred direction and remained below 0.01 in the opposite direction; all remaining cells were classified as central. For Fig. 5a, the mean firing of each cell group was computed by first averaging across 32 trajectories with the same speed but different initial positions, and then averaging across all cells within the group. Although only clockwise trajectories are shown, equivalent results were obtained for counterclockwise motion.

Figures 6b,c and d were generated from the unstructured task, with a time stimulus presented only at *t* = 0. Panel (b) was generated like figure 3c. In the case of panel (c), we calculated the normalized mean activity of each cell across all trials, and then sorted all time cells according to the time with highest mean firing. Panel (d) was generated by calculating two-dimensional space-time firing maps.

### 4.8 Attractor speed estimation

To calculate the speed inside the attractor for models in figures 3, 5 and 6, we simulated trajectories starting from random positions, with fixed speed and direction. Then, to control for the fact that different cells have different activity ranges, we computed a normalized activity vector **x̃***^t^* by dividing each component by its maximum value across all episodes and time steps. Then, we computed the attractor speed at time *t* as the norm of the difference between normalized activity vectors **x̃** at two consecutive time steps: ||**x̃***^t^*^+1^ − **x̃***^t^*||. We averaged the estimated attractor speed across all times and 16 trials with the same speed but different initial positions.

### 4.9 Model perturbations

To test the proposed brake role of the central-cell population, we weakened the connections from the speed input to these cells. Because the original speed weights included both positive and negative values, direct scaling would simultaneously reduce inhibitory and excitatory influences. We therefore first subtracted the maximum speed-to-central weight from all speed, direction, and visual input weights projecting to the recurrent layer. Since the Softmax activation is invariant to adding the same constant to all its inputs, this transformation does not alter the network dynamics but makes all speed-to-central weights non-positive. We then progressively multiplied the transformed speed-to-central weights by decreasing scaling factors and evaluated the model after each perturbation. This manipulation therefore uniformly reduced the suppressive effect of speed on the central-cell population.

After each perturbation, we calculated the angle estimated by the output 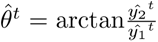 in trajectories with fixed speed and direction, for both the perturbed and unperturbed models.

*θ̂* ∈ [0, 2*π*), trajectories exhibited artificial discontinuities whenever the estimated angle crossed the 2*π* → 0 boundary. To obtain a continuous representation of the decoded angle, the angular trajectories were phase-unwrapped by adding 2*π* whenever the difference between two consecutive estimates exceeded −*π*, corresponding to a wrap-around from 2*π* to 0. This transformation removes discontinuities introduced by the angular representation without altering the underlying trajectory.

Then, linear regressions were fitted between the estimated angle for the unperturbed and the perturbed models inside each trial. Finally, we averaged the slopes of all trials for every perturbation.

### 4.10 Simplified recurrent network with connectivity motifs from the trained RNN

To generate the recurrent matrix, we defined each column as a gaussian function, 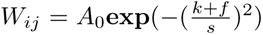 where *k* is the circular distance between the *i*-th and *j*-th cell. Then *f*, the shift from the diagonal, was chosen as 4.5 for CW cells, -4.5 for CCW cells, and 0 for central cells. The final matrix was generated by alternating values of these phases. The speed input weight was set to −*µ* for central cells only, and 0 for directional cells. *µ* was varied across models. Direction input weights were set to −0.2, but only for cells that encoded the opposite direction.

We defined positions in the speed sorted rate maps by setting *p^t^*^=0^ = 0 and computing *p^t^*^+1^ = *p^t^* + *v dt*. In all cases this initial position is corresponded by equally setting the activations of the first central cell to 1.

For each animal speed, we computed the attractor speed as described for the trained model and averaged attractor speed values across time steps and 32 trials initiated from different cells.

### 4.11 Recurrent neural networks trained to encode space and time

The timing task consisted on the presentation of a cue at random times, from which the model had to estimate the elapsed time. We randomly sample time intervals from a uniform distribution in the range [2*, τ*] with *τ* being the total number of timesteps in the trial. Then, we constructed the sequence of cues according to those intervals. Every trial started with a cue presentation at *t* = 0. The network architecture was identical to that of the space-only model, and the loss function was

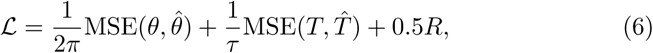

where *T* is the target elapsed time and *T̂* is the model’s estimate.

All model parameters were identical to those of the space model, shown in table 4.5.2, except that *τ_T_*= 20*, β_T_* = 15.

**Table 1:** Training and Network hyper-parameters for the spatial and the time-spatial model.

| Parameter | Description | Value |
| --- | --- | --- |
| **Task** |  |  |
| $v$ | Animal speed in training | $\sim N(0, 1)$ |
| $P(v = 0)$ | Probability of $v = 0$ | 0.1 |
| $\tau_S$ | Number of simulation time steps (space model) | 40 |
| $\tau_T$ | Number of simulation time steps (space and time model) | 20 |
| $dt$ | Duration of timestep | 0.2 s |
| $N_{vis}$ | Presentations of visual stimuli | 5 |
| **Network** |  |  |
| $N$ | Number of cells | 128 |
| $\beta_S$ | Softmax inverse temperature (space model) | 20 |
| $\beta_T$ | Softmax inverse temperature (time model) | 15 |
| **Training** |  |  |
| $B$ | Batch size in training | 128 |
|  | Learning rate | 1e-3 |
| $\epsilon$ | Pre-activation noise | $\sim N(0, 0.01)$ |

## 5 Data availability

The data generated during this study are available from the corresponding author upon reasonable request. No experimental datasets were used in this study.

## 6 Code availability

All code that supports the findings of this study is available at https://github.com/fszmidt/PlaceTimeShifts

## 7 Acknowledgements

F.S. is supported by a doctoral fellowship from the Consejo Nacional de Investigaciones Cientìficas y Técnicas (CONICET), Argentina.

## 8 Author contributions

C.J.M. conceived the study and supervised the project. F.S. developed and implemented the recurrent neural network models and performed the computational analyses. C.J.M. developed the line-attractor model. F.S. and C.J.M. interpreted the results, wrote the manuscript, and approved the final version.

## 10 Additional information

Correspondence and requests should be addressed to Camilo J. Mininni

### 1 Supplemental information

**Figure S1:**
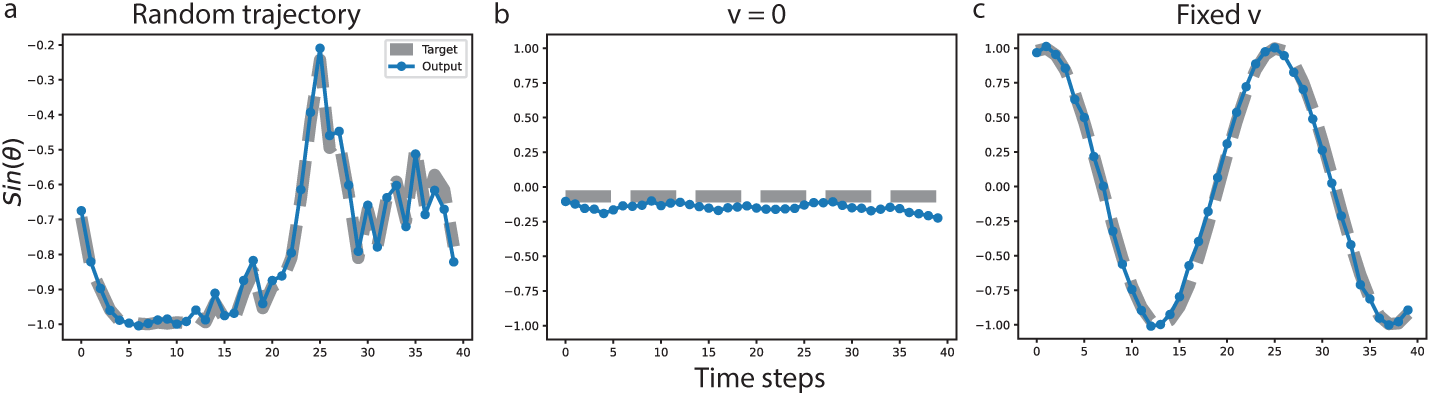
Output examples of the RNN trained only to solve a path-integration task. Left: Example trajectory as used in training. Center: Example trajectory with zero velocity, showing the stability of the attractor. Right: Example trajectory with fixed speed, as used to generate Figure 3d

**Figure S2:**
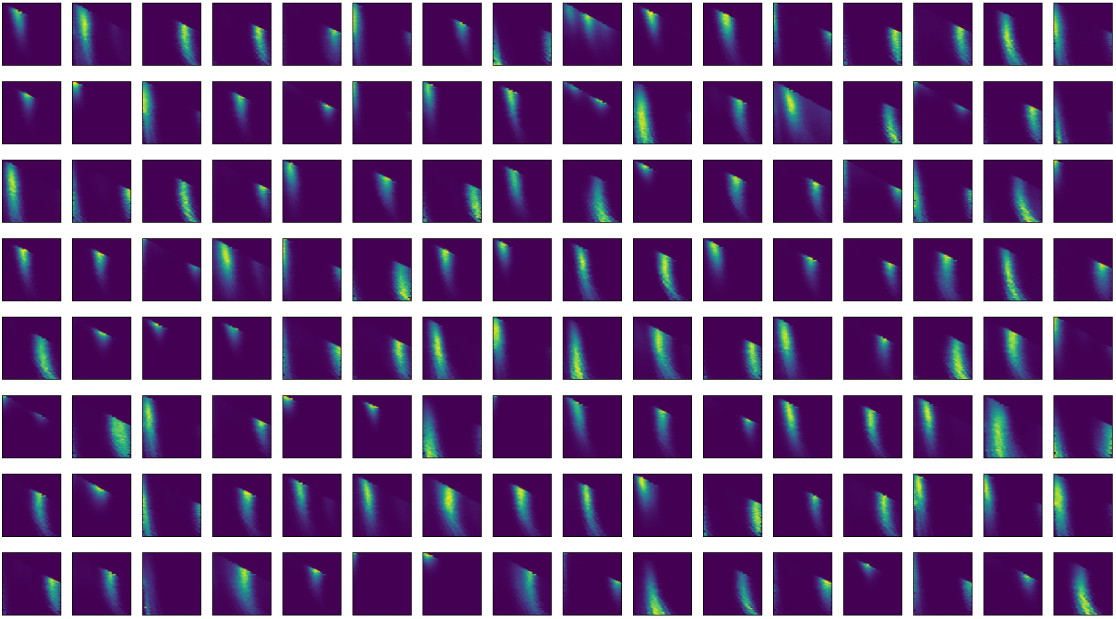
Speed-ordered spatial rate maps for all cells in the RNN trained in the spatial task during the structured task.

**Figure S3:**
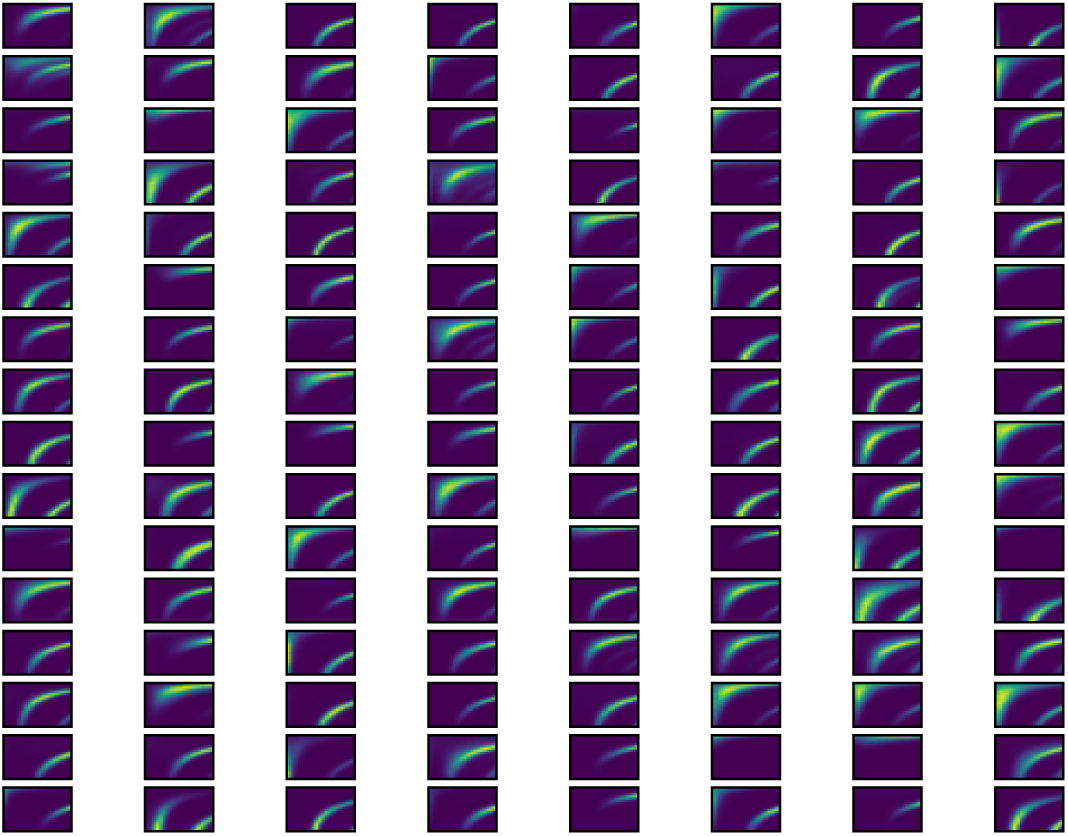
Speed-ordered temporal rate maps for all cells in the RNN trained in the spatial task during the structured task.

**Figure S4:**
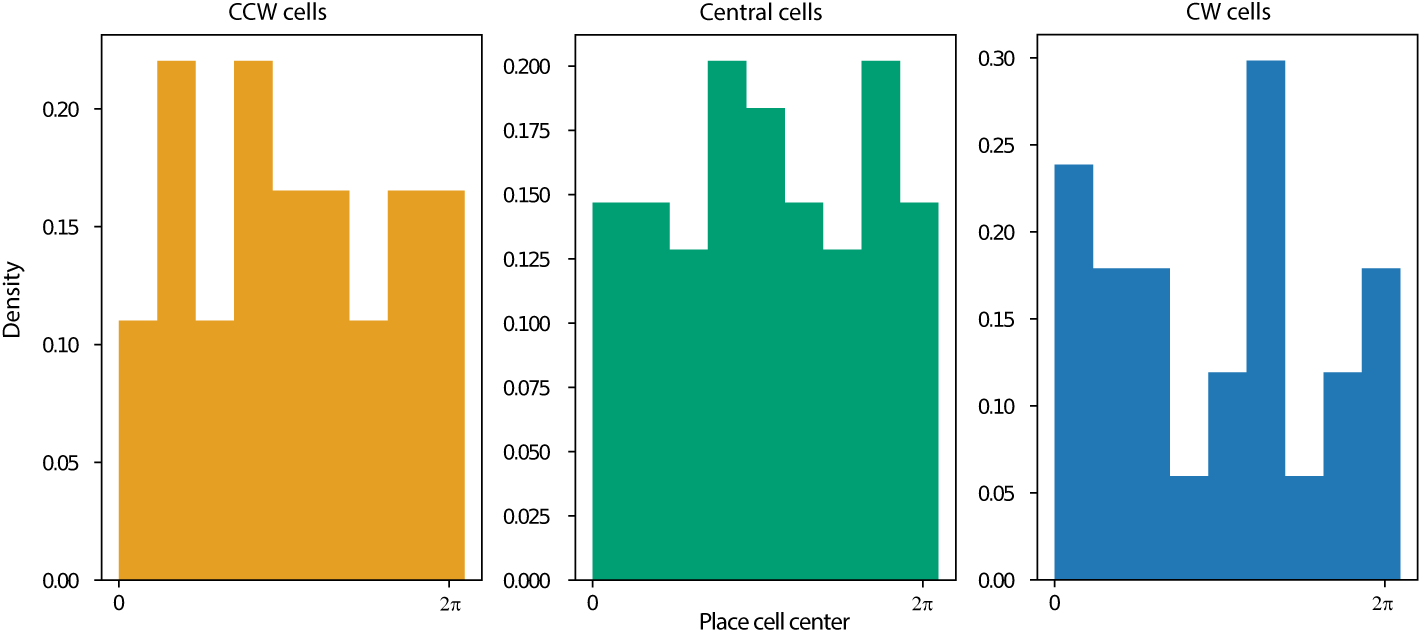
Position selectivity distribution for each population in the model trained for the spatial task.

**Figure S5:**
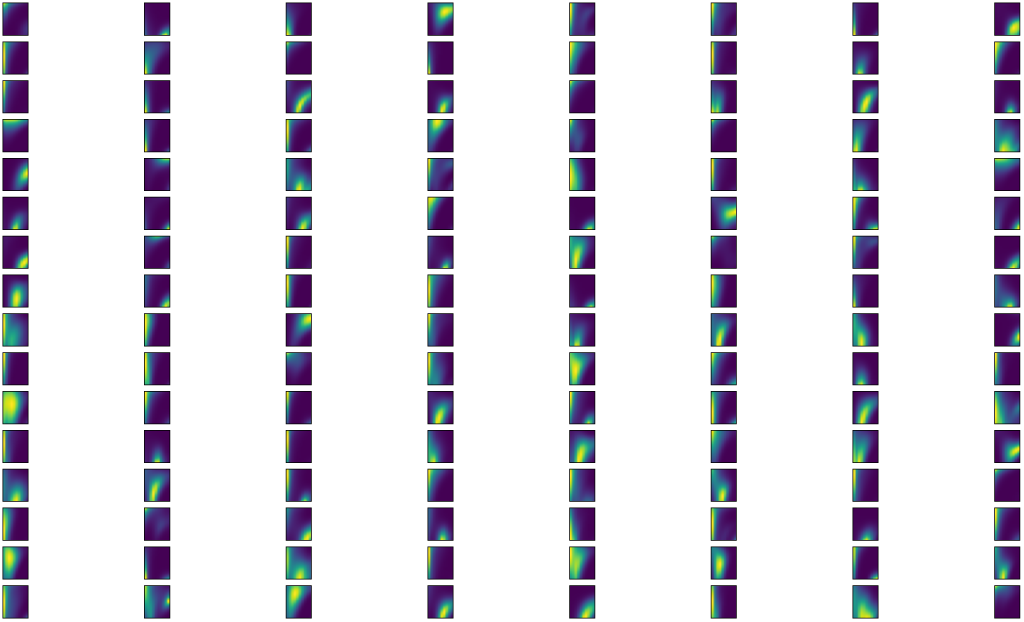
Speed-ordered spatial rate maps for all cells in the RNN trained in the spatial and temporal task during the structured task.

**Figure S6:**
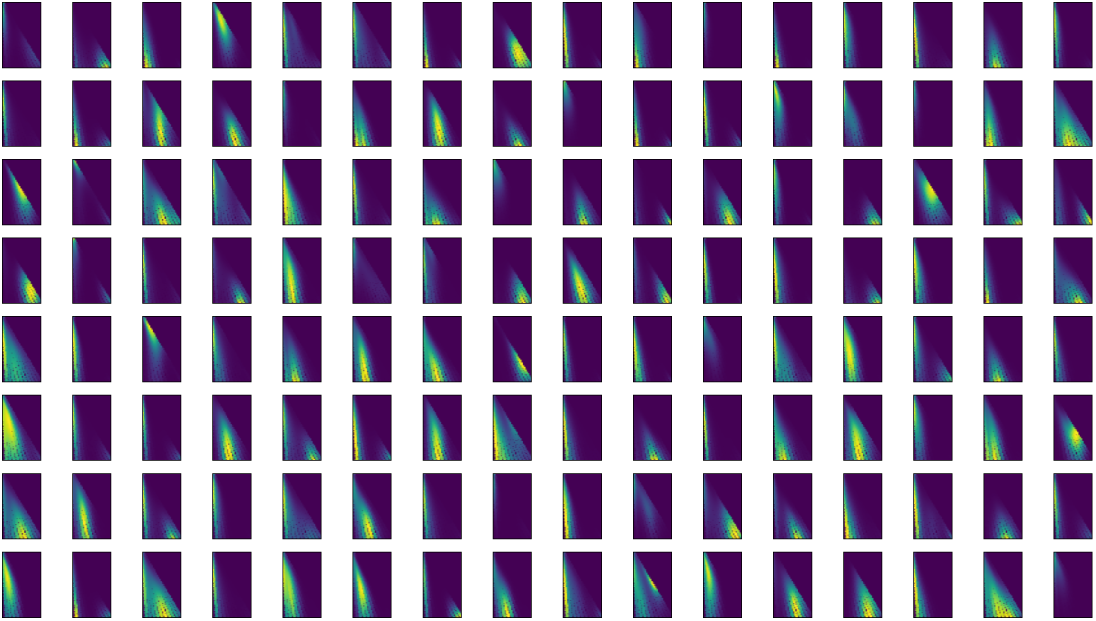
Speed-ordered temporal rate maps for all cells in the RNN jointly trained in the spatial and temporal task during the structured task.

**Figure S7:**
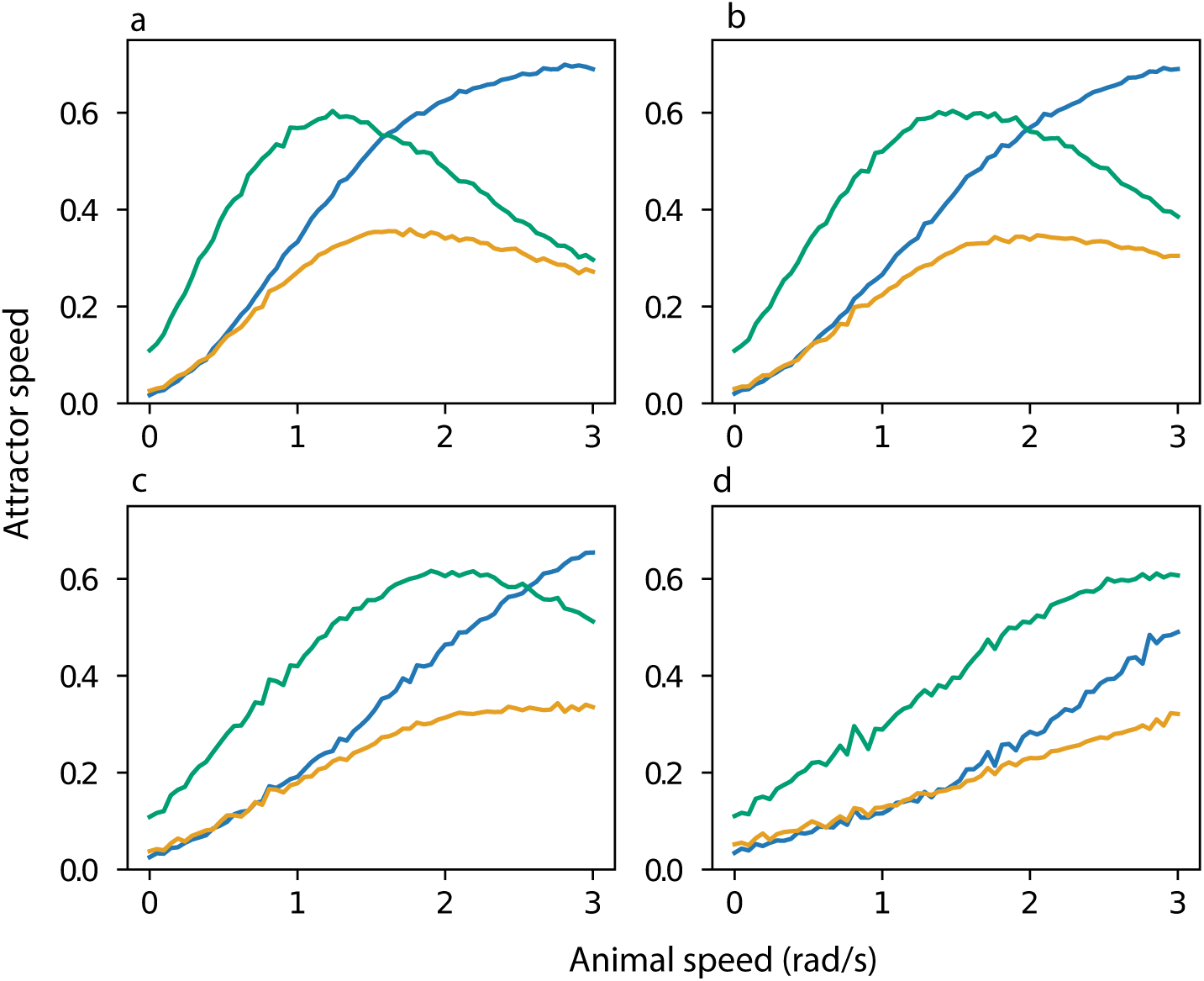
Normalized attractor speed for each subpopulation in the model trained for the spatial task (a) and after the three perturbations (c-d)

**Figure S8:**
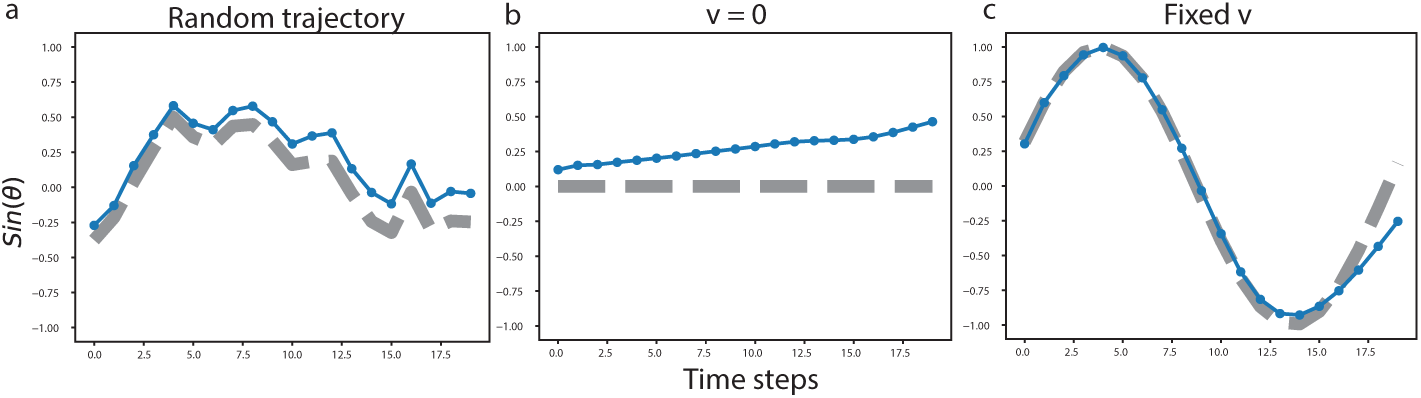
Same as S1 but for the model trained to jointly solve a path-integration task, and a timing task. In all cases the time cue was only presented at t = 0.

**Figure S9:**
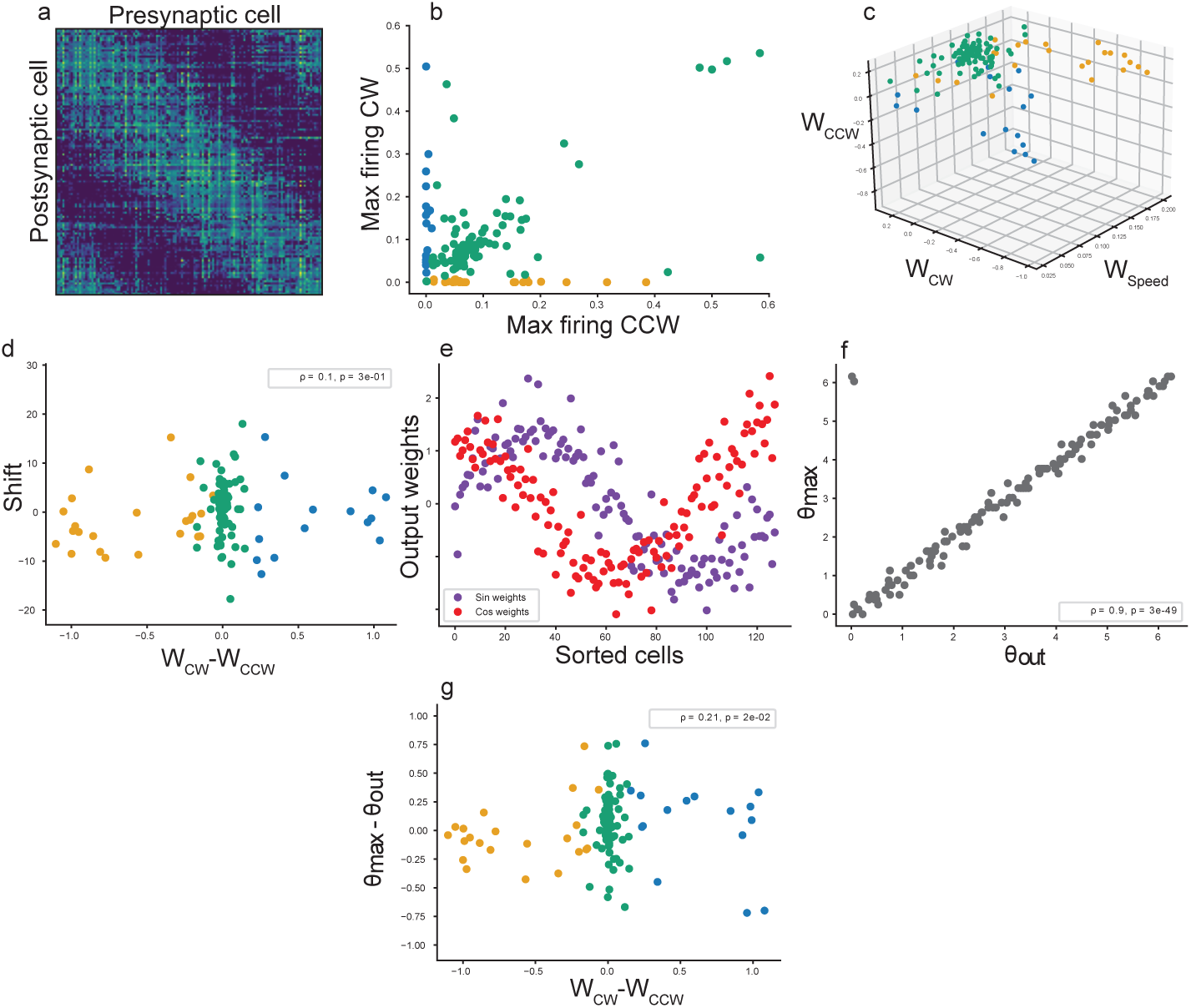
**An RNN trained to solve a path-integration task and a timing task encodes space like a CBAN**. **a** Recurrent weight matrix sorted by each cell’s preferred position *θ_max_*. Columns correspond to outgoing weight vectors and rows to incoming weight vectors. Thus, matrix entry (*i, j*) represents the weight of the connection from the neuron in column *j* to the neuron in row *i*. **b** Maximum firing of each cell in trajectories with fixed directions; cells are classified as direction-selective if they have a maximum firing *>* 0.01 in the preferred direction and < 0.01 in the opposite direction, and as central cells otherwise. **c** Direction and speed input weights for all cells. **d** Recurrent shift, defined as the distance between the center of mass of each column and the matrix diagonal, plotted against the difference between CW and CCW input weights. The recurrent shift is not significantly correlated (Spearman’s *ρ* = −0.1, *p* = 0.17). **e** Output weights *O*_1_, *O*_2_ corresponding to targets *sin*(*θ*), *cos*(*θ*), for all units in the network, sorted by the angle of maximum firing *θ_max_*. **f** The preferred output angle 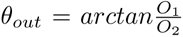 closely matches *θ_max_* for each cell. **g** Cells with stronger inputs favoring the CW or CCW direction have preferred output angles shifted ahead or behind their preferred firing position.

**Figure S10:**
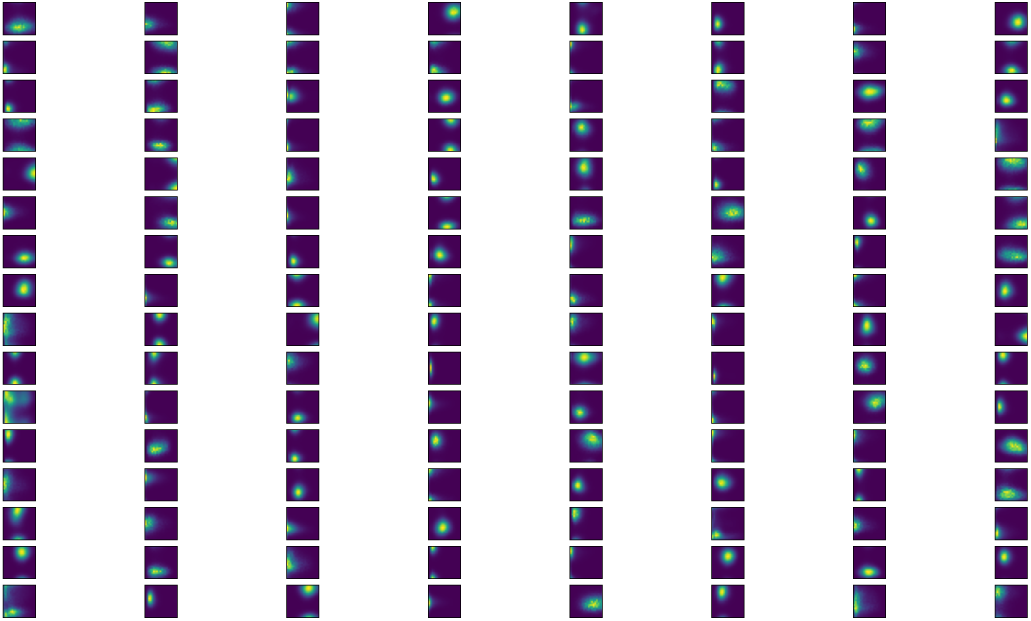
Position-time rate maps for all cells in the RNN jointly trained in the spatial and temporal task during the structured task.

